# Transcriptional regulation in the absence of Inositol Trisphosphate Receptor Calcium Signaling

**DOI:** 10.1101/2024.04.16.589553

**Authors:** Michael Young, David M. Booth, David Smith, Marco Tigano, Gyӧrgy Hajnόczky, Suresh K. Joseph

## Abstract

The activation of IP_3_ receptor (IP_3_R) Ca^2+^ channels generates agonist-mediated Ca^2+^ signals that regulate a wide range of biological processes. It is therefore surprising that CRISPR induced loss of all three IP_3_R isoforms (TKO) in HEK293 and HeLa cell lines yields cells that can survive, grow and divide, albeit more slowly than wild-type cells. In an effort to understand the adaptive mechanisms involved, we have examined the activity of key Ca^2+^ dependent transcription factors (NFAT, CREB, AP-1 and NFκb) and signaling pathways using luciferase-reporter assays, phosphoprotein immunoblots and whole genome transcriptomic studies. In addition the role of protein kinase C (PKC) was investigated with inhibitors and siRNA knockdown. The data showed that agonist-mediated NFAT activation was lost but CREB activation was maintained in IP_3_R TKO cells. Under base-line conditions transcriptome analysis indicated the differential expression (DEG) of 828 and 311 genes in IP_3_R TKO HEK293 or HeLa cells, respectively, with only 18 genes being in common. In summary three main adaptations in TKO cells are identified in this study: 1) increased basal activity of NFAT, CREB, AP-1 and NFκb; 2) an increased reliance on Ca^2+^-insensitive PKC isoforms; and 3) increased production of reactive oxygen species and upregulation of antioxidant defense enzymes. We suggest that whereas wild-type cells rely on a Ca^2+^ and DAG signal to respond to stimuli, the TKO cells utilize the adaptations to allow key signaling pathways (e.g. PKC, Ras/MAPK, CREB) to transition to the activated state using a DAG signal alone.

## INTRODUCTION

Calcium is viewed as a vital signaling molecule that regulates many important physiological processes including growth, cell division, motility, gene expression and metabolism (1). Agonist-mediated Ca^2+^ signals are evoked as a result of the formation of inositol 1,4,5-trisphosphate (IP_3_) which opens IP_3_R channels in endoplasmic reticulum membranes. The accompanying depletion of intracellular stores secondarily activates the STIM/Orai pathways of Ca^2+^ entry across the plasma membrane. Three IP_3_R isoforms are expressed in cells and several labs have successfully genetically deleted all three isoforms to generate IP_3_R triple knockout cells (TKO). This includes chicken DT40 cells (2), mouse T and B lymphocytes (3,4), HEK293 cells (5,6), HeLa cells (7), mouse embryonic stem cells (8) and human induced pluripotent stem cells (9). All these model systems completely lack agonist-mediated Ca^2+^ signals. In view of the proposed central role of Ca^2+^ signaling, it is surprising that these cells display a somewhat mild phenotype and continue to grow and divide, in some cases more slowly (6,10–12), and in some cases at rates indistinguishable from wild type cells (8,9). This suggests that TKO cells have adapted to the loss of Ca^2+^ signaling. However, the compensatory mechanisms that enable these cells to maintain homeostasis and reconfigure their transcriptional landscape in the absence of Ca^2+^ have not been investigated.

Many transcriptional regulators are modulated by Ca^2+^ signals. Early studies showed that Ca^2+^ signals rapidly induced expression of the proto-oncogene c-fos upon stimulation of receptors for acetylcholine in PC12 cells (13) or growth factors in quiescent fibroblasts (14). Both NFAT and CREB are also activated by increased Ca^2+^ signaling (15–17). AP-1, which are transcription factors composed of dimers of fos and jun family members, are also regulated by Ca^2+^ (18,19). Multiple signaling pathways are involved in the activation of Ca^2+^-sensitive transcription factors including protein kinase C (PKC), RAS/MAPK and Jnk. In the present study, we have used luciferase reporter assays to measure the activity of transcription factors and biochemical assays to monitor signaling pathways. We used TKO cells from two different human cancer cell lines (HEK293 and HeLa) to identify the Ca^2+^-dependent transcriptional pathways that have been inactivated or that continue to function. We have also examined the global expression of genes by RNAseq in both models. Although there are some differences in the behavior of the two cell lines, our main findings are that TKO cells show an increased baseline activity of NFAT, CREB and AP-1, an increased reliance on Ca^2+^-insensitive PKC isoforms, and a hitherto unrecognized alteration in the handling of ROS and redox poise. All of these changes may contribute to the ability of these cells to grow and divide in the absence of Ca^2+^ signaling.

## RESULTS

### Higher basal activity and altered Ca^2+^ sensitivity of NFAT in HEK293 TKO cells

To investigate the impact of Ca^2+^ signal loss on transcription, we began by looking at the activation of the well-established Ca^2+^-dependent transcription factor NFAT using a luciferase reporter assay. We expressed the effect of treatments as a fold-activation over the baseline of WT cells, after normalization for transfection efficiency using Renilla luciferase activity. However, a 50-70% increase in baseline NFAT activity was noted in both HEK293 TKO cells and HeLa TKO cells (**Figure 1A&C**). HEK293 cells express endogenous M3 muscarinic receptors (20) and stimulation with carbachol (Cch) caused a 3.5-fold increase in NFAT activity in WT cells, but had no further stimulatory effect above the baseline of TKO cells (Fig 1A). Activation of endogenous histamine receptors in HeLa cells showed a 2-fold activation of NFAT in WT cells which is lost in TKO cells (Fig 1B). These observation are in line with published data showing loss of NFAT stimulation by B-cell receptor activation in DT40 TKO cells (2), and reduced NFAT activation in HEK293A cells after siRNA knockdown of type 2 IP3Rs (21). As a control to verify that NFAT was still responsive to Ca^2+^ in TKO cells, we tested the effect of the Ca^2+^ ionophore ionomycin. The ionophore had comparable effects to agonists in both WT HEK293 and HeLa cells. However, ionophore elicited an unexpectedly large increase (∼13-fold activation) in NFAT activity in the HEK293 TKO cells (Fig 1A). An increased NFAT response to thapsigargin-induced elevation of cytosolic Ca^2+^ was also observed in HEK293 TKO cells (Fig 1B). Previous studies have shown equivalent responses of [Ca^2+^]c to ionomycin and thapsigargin in HEK293 TKO cells (6,1). The hypersensitivity to ionomycin was not observed in the HeLa TKO model. These results suggest that one adaptation to a loss of Ca^2+^ signaling in HEK293 TKO cells is an increased Ca^2+^ sensitivity of the calcineurin (CaN) -mediated dephosphorylation of NFAT. To test for the involvement of CaN we pretreated the cells with cyclosporine A which blocked the ionomycin responses in both WT and TKO HEK 293 cells and eliminated the baseline differences seen in the TKO cells (Fig 1A). The enhanced sensitivity of TKO cells to TG was also inhibited by cyclosporine A (Fig 1B). Many possible mechanisms may underlie the altered Ca^2+^ sensitivity of CaN regulation in HEK293 TKO cells, including a slower export of NFAT from the nucleus or altered levels of endogenous CaN inhibitors such as RCAN1 (22). We found decreased levels of RCAN1 in HEK293 TKO cells but not in HeLa TKO cells (Fig 1D), which would be consistent with supersensitivity to ionomycin being confined to HEK293 cells. Another reason for the lack of Ca^2+^ hypersensitivity could be related to HeLa cells having a different complement of NFAT isoforms than HEK293 cells (23), since each isoform also shows differential regulation and kinetics of nuclear translocation (24,25).

**Figure 1.**
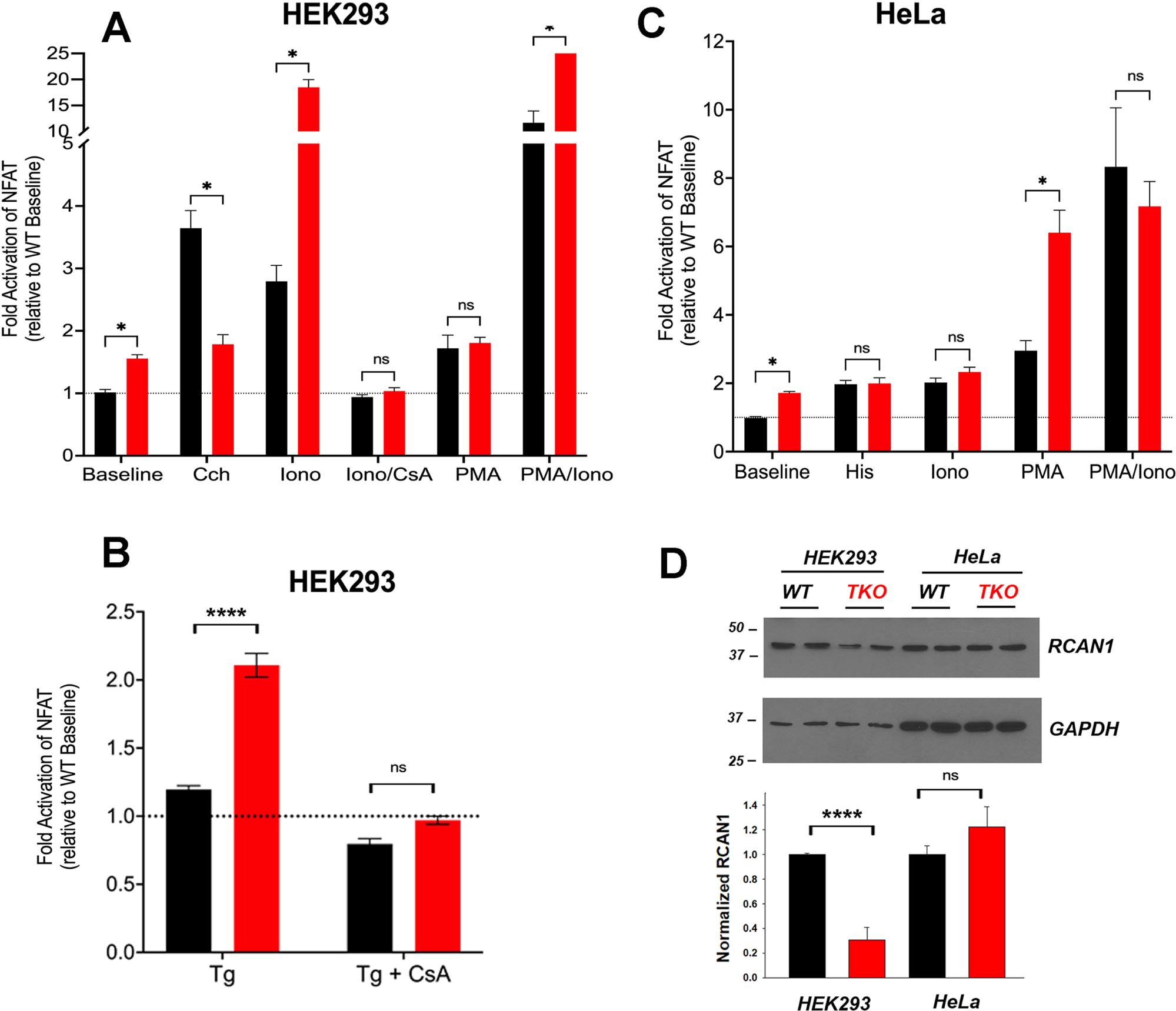
NFAT activity in wild-type and IP_3_R TKO cells. ***Panel A*** HEK293 WT (black) and TKO (red) cell lines were co-transfected with fire-fly NFAT-luciferase and *Renilla* luciferase reporter vectors. After 24h, cells were treated for a further 4h with either 25μM carbachol (Cch), Ionomycin (Iono) or 100nM PMA. When used, 0.5μM cyclosporine A (CsA) was preincubated for 30min. Lysates were assayed for luciferase activity as given in ‘Experimental Procedures’. All fire-fly/Renilla luciferase ratios were normalized to the value obtained for WT cells at baseline. Statistical significance * p<0.05; ****p<0.0001; ns=not significant. ***Panel B*** HEK293 cells were treated with 0.5µM Thapisgargin (Tg) in the presence or absence of cyclosporin A (CsA) pretreatment as in Panel A. ***Panel C*** HeLa WT (black) and TKO (red) cell lines were treated with 25μM Histamine (His) and other reagents as described in Panel A. ***Panel D*** HEK293 and HeLa WT (black) and TKO (red) cell lines were immunoblotted for RCAN1. In the lower panel RCAN1 levels were normalized to GAPDH and quantitated. The data shown are the mean ± S.E.M.of 3 independent experiments.

Many NFAT-dependent target genes are also stimulated by the AP-1 transcription factor, which in turn is activated by PKC signaling (26). The co-operative binding of NFAT and AP-1 to adjacent promoter sites results in synergistic activation of NFAT gene targets when cells are simultaneously stimulated by Ca^2+^ and PKC activation (27). A pan PKC activator, phorbol 12-myristate 13-acetate (PMA), caused a ∼70% activation of NFAT in the HEK293 WT or TKO cell line (Fig 1A). Combination treatment of PMA and Ionomycin elicited synergistic responses in WT and TKO cells. However, the PMA/Ionomycin treatment caused a larger increase in the TKO cells (45-fold versus 12-fold in WT cells). In HeLa cell lines, PMA caused a larger NFAT activation in TKO cells (Fig 1C). A synergistic activation by PMA/Ionomycin was seen in the HeLa WT line but not in TKO cells which were already maximally activated by PMA alone (Fig 1C).

### Higher basal activity, altered Ca^2+^ sensitivity and increased reliance on Ca^2+^ independent PKC for CREB activation in TKO cells

Next, we investigated cAMP response element binding protein (CREB), another well studied Ca^2+^-sensitive transcription factor (28). Ca^2+^-modulation of CREB is exerted through phosphorylation by specific isoforms of two protein kinase families, Ca^2+^/calmodulin-dependent protein kinase (CaMK) and protein kinase C (PKC) (28,29). Using a CRE-luciferase reporter assay to monitor that activity of CREB, we observed baseline increases in both HEK293 and HeLa TKO cells (**Figure 2A & B**), but unlike NFAT, CREB luciferase activity was still stimulated by agonists in TKO cells of both cell types. The addition of ionomycin to the WT cells had a smaller effect than agonist on CREB-luciferase activity. However, ionomycin promoted a marked increase in CREB reporter activity (>10-fold) in HEK293 TKO cells. Cyclosporine A blocked the effects of ionomycin and eliminated the baseline increase in CREB reporter activity seen in the TKO cells (Fig 1A). The supersensitivity to ionophore was not observed in HeLa TKO cells which were actually less sensitive to ionophore compared to WT cells (Fig 2B). The selective effect of ionomycin resembles that seen with NFAT luciferase in HEK293 cells (Fig 1A) and may have the same underlying mechanism. It is known that CaN can activate CREB via dephosphorylation and nuclear translocation of CRTC2, a transcriptional coactivator of CREB (30). Thus, decreased levels of RCAN1 could play the same role in altering the Ca^2+^ responsiveness of CREB in a manner similar to NFAT. The different ionophore response in HeLa cells may be due to the lack of RCAN1 changes, but it should be noted that the regulation of CRTC2 is also different in HeLa cells since they lack an upstream kinase (LKB1) needed to maintain phosphorylation of CRTC2 (31). In a previously published study on HEK293A cells it was shown that optimal responses of CREB to Ca^2+^ mobilizing hormone requires the addition of vasoactive intestinal peptide (VIP), a cAMP-elevating stimulus (21). Treatment with VIP alone or in combination with ionophore caused a very large activation of CREB in both cell lines which was not significantly different in the TKO cells (Fig 2A & B).

**Figure 2.**
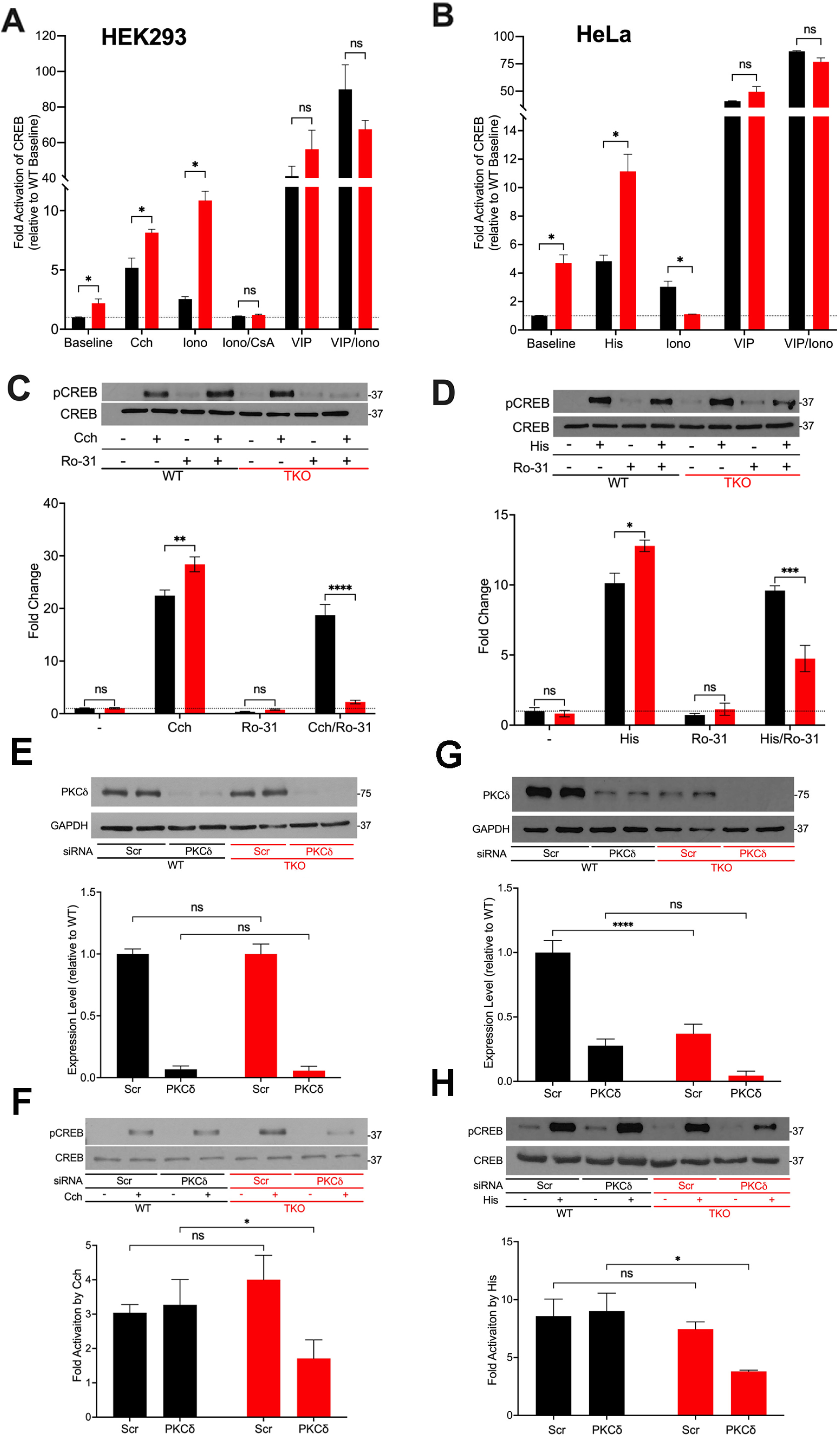
CREB activity in wild-type and IP_3_R TKO cells. ***Panel A*** HEK293 WT (black) and TKO (red) cell lines were co-transfected with fire-fly CRE-luciferase and *Renilla* luciferase reporter vectors. Treatment conditions and data analysis were as described for Fig 1A. Vasoactive intestinal peptide (VIP) was used at 0.5μM. ***Panel B*** HeLa WT (black) and TKO (red) cell lines were treated with reagents as described in Panel A except 25μM Histamine (His) was used as agonist. ***Panel C*** Representative immunoblots for pCREB (ser-133) and total CREB in HEK293 WT and TKO lysates are shown. Cells were treated for 10min with 25µM carbachol (Cch), 1μM Ro-31-8220 (Ro-31), a pan-PKC inhibitor, or both. In the lower panel p-CREB data was expressed as a ratio of total CREB and normalized to baseline WT values. ***Panel D*** Representative immunoblots and quantitation of pCREB in HeLa WT and TKO lysates are shown. Conditions were the same as used for Panel C except for the use of 25μM Histamine (His) as agonist. ***Panel E*** HEK293 WT and TKO cells were transfected with scrambled siRNA (Scr) or siRNA for PKCδ for 48h and expression levels of PKCδ was measured by immunoblotting using GAPDH as a loading control. Expression was normalized to WT levels and the data shown are the mean ± S.E.M.of 3 independent experiments. ***Panel F*** HEK293 WT and TKO cells were transfected with siRNA as described in Panel E and then stimulated for 10min with 25μM carbachol (Cch). Changes in p-CREB and total CREB were measured by immunoblotting. Fold activation by Cch was quantitated in 3 experiments in the lower panel. ***Panel G & H*** Conditions were the same as described for Panels E & F except HeLa WT and TKO cells were used and were stimulated with 25μM Histamine (His) as agonist.

A primary phosphorylation site in CREB is at ser-133 which enhances recruitment of the co-activator CBP/p300 and increases transcriptional activity (32). To confirm the data obtained with CREB reporter assays, we immunoblotted for p-CREB (ser-133) after 10min treatment with an agonist in HEK293 (Fig 2C) and HeLa (Fig 2D) cells. Agonist stimulation resulted in increased CREB phosphorylation in both WT and TKO cells, with agonist effects being significantly larger in the TKO cell lines (Fig 2C & D). To investigate the role of PKC in mediating CREB phosphorylation we used a pan-PKC inhibitor, Ro-31-8220 (33). The inhibitor did not block agonist-stimulated pCREB in WT HEK293 or HeLa cells but it suppressed phosphorylation in both TKO cell lines (Fig 2C & D). Overall, the results show that agonist mediated CREB activity and phosphorylation is maintained in TKO cell lines and that TKO cells are more reliant on PKC signaling for CREB phosphorylation at Ser133. In order to identify the isoform of PKC that may be involved in maintaining CREB activation in TKO cells we used an siRNA knockdown approach. The obvious candidates are the novel Ca^2+^-insensitive isoforms, of which PKCδ and PKCε are both expressed in HEK293 and HeLa cells (34,35). HEK293 WT and TKO cells expressed similar basal levels of total PKCδ and treatment with siRNA resulted in >90% knockdown in both cell lines. (Fig 2E). Knockdown of PKCδ had no effect on agonist stimulated CREB phosphorylation in HEK293 WT cells but caused a 50% reduction of pCREB in TKO cells (Fig 2F). In HeLa cells, expression levels of basal PKCδ in TKO cells was only 25% of that seen in WT cells (Fig 2G). Treatment with siRNA reduced PKCδ in HeLa WT and TKO cells but resulted in significantly reduced pCREB only in TKO cells (Fig 2H). The poor quality of our PKCε antibodies precluded further investigation of the contribution of this isoform. Overall, these data indicate that CREB activation is maintained in TKO cells, in part, by an increased reliance on Ca^2+^-insensitive PKC isoforms.

### AP-1 is constitutively active in TKO cells

AP-1 is a transcription factor composed of homo- and heterodimers of members of the fos, jun, Maf and ATF families. These proteins are categorized as “immediate early genes” that are normally present at low amounts in quiescent cells and are induced in response to activation of signaling pathways, such as Ca^2+^ mobilization or the activation of PKC. Initially, we examined the activity of AP-1 in WT and TKO cells using luciferase promoter assays (**Figure 3**). The results show that agonists were able to stimulate AP-1 activity in the WT HEK293 (Fig 3A) and HeLa cells (Fig 3B) by ∼2-fold. However, the TKO cell lines already had elevated baseline AP-1 activity, which could not be further increased by agonists. AP-1 activity in HEK293 cells was not responsive to ionomycin but was stimulated by PMA. This is in line with the known sensitivity of AP-1 to activation by PKC (36). Although PMA treatment of TKO cells gave an apparent smaller fold activation in both cell types, the absolute levels of AP-1 activity were comparable when the elevated baselines were taken into account. The combined treatment with PMA and ionomycin produced a synergistic activation in both WT and TKO HEK293 cells but not in HeLa cells.

**Figure 3.**
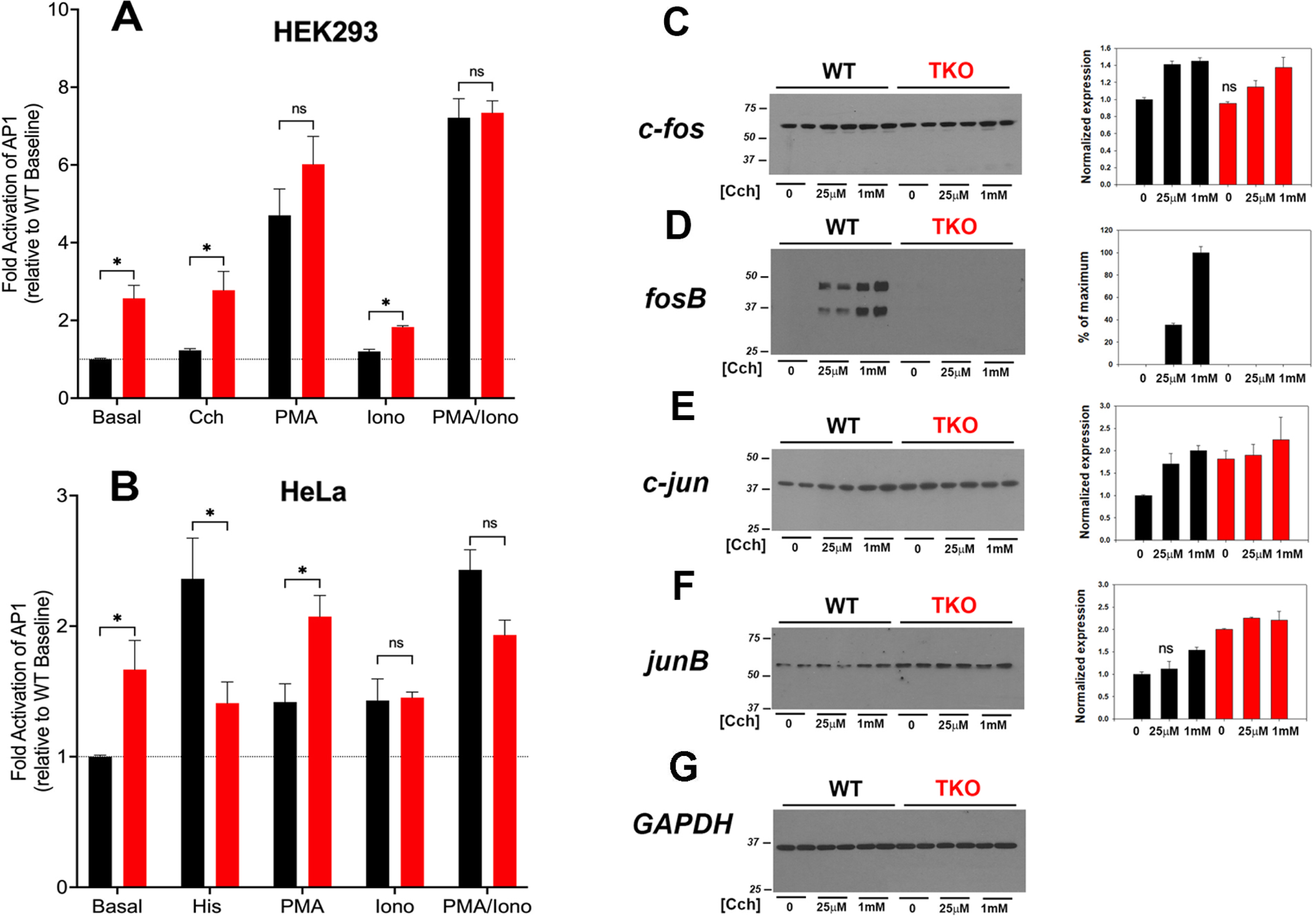
AP-1 activity in wild-type and IP_3_R TKO cells. ***Panel A*** HEK293 WT (black) and TKO (red) cell lines were co-transfected with fire-fly AP1-luciferase and *Renilla* luciferase reporter vectors. Treatment conditions and data analysis were as described for Fig 1A. ***Panel B*** HeLa WT (black) and TKO (red) cell lines were treated with reagents as described in Panel A except 25μM Histamine (His) was used as agonist. ***Panel C*** Lysates were prepared from HEK293 WT and TKO cells treated with the indicated concentrations of Cch for 4h. Samples were immunoblotted for c-fos and quantitated using GAPDH as a loading control. Data were normalized to the WT unstimulated values. ***Panel D*** fosB; ***Panel E*** c-jun; ***Panel F*** junB. The data shown are the mean ± S.E.M. of 3 independent experiments. All data are statistically significant from WT unstimulated values at p<0.05 with only the data not significantly different marked “ns”.

The protein expression levels of four AP-1 members (c-fos, fosB, c-jun, junB) were examined by immunoblotting in cell lysates of control and agonist-stimulated HEK293 WT and TKO cells (Fig 3C-G). Significant levels of c-fos, c-jun and junB were detected at baseline even in serum-deprived HEK293 cells. In the case of c-fos, basal levels were comparable in WT and TKO cells and carbachol treatment increased c-fos similarly in both cell lines, although higher concentrations of carbachol were required in TKO cells (Fig 3C). By contrast, baseline fosB was not detectable in WT or TKO cells and was increased by carbachol treatment only in WT cells (Fig 3D). Two bands were detected for fosB with both bands showing the same changes. Based on its molecular weight, the lower band likely corresponds to the molecular weight of the established splice variant fosB2 (also known as delta fosB)(37). c-Jun (Fig 3E) and JunB (Fig3F) behaved similarly, with protein levels increasing in response to carbachol in WT cells. However, baseline levels of both proteins were elevated in TKO cells and could not be further stimulated by carbachol treatment. Changes in the levels of these AP-1 proteins in WT and TKO HeLa cells are shown in Fig S1. c-Fos and JunB in HeLa cells behaved in a manner similar to c-jun and JunB in HEK cells. Notably, baseline levels were increased in TKO cells and could not be further stimulated by histamine. The histamine response of c-jun was decreased in TKO cells and FosB was not expressed in HeLa cells under any experimental conditions (not shown). The results indicate that some members of the AP-1 family are more highly expressed at baseline in TKO cells and this could underlie the constitutive activation of AP-1 activity seen under these conditions.

### MAP kinase and NFκb activities are maintained in TKO cells

The activation of ERK1/2, p38 and Jnk MAP kinase cascades lie upstream of the AP-1 and CREB pathways. The consensus of studies suggests that while Ca^2+^ signals are not essential, they can regulate these pathways. For example Ca^2+^ elevation in primary cortical neurons (38,39), fibroblasts (40) and HeLa cells (41) activates the Ras/Raf/ERK1/2 pathway. Similarly, there have been many studies which report the involvement of a Ca^2+^ signal in the activation of p38 and Jnk (42) pathways. We have examined the carbachol-mediated activation of Jnk and ERK1/2 in HEK293 TKO cells (Fig S2). In the case of Jnk, a 54kDa species was phosphorylated in response to carbachol stimulation in both WT and TKO cells (Fig S2A-C). However, a higher baseline level and more rapid kinetics of p-Jnk formation were evident in the TKO cells. Similarly, the ERK1/2 pathway was activated robustly by carbachol in both WT and TKO cells (Fig S2D&E).

Multiple mechanisms have been identified for the Ca^2+^ regulation of NFκb activity including activation of PKC (43). A NFκb luciferase promoter showed a higher basal activity in HEK293 TKO cells but was insensitive to carbachol or ionophore in either WT or TKO cells (Fig S2F&G). As expected PMA was a potent stimulus for NFκb promoter activity but the combination of ionophore and PMA was less effective, particularly in TKO cells (Fig S2). HeLa WT and TKO cells showed several differences from HEK293 cells including similar baseline activity and less pronounced sensitivity to PMA (Fig S2G). In contrast to lymphocytes, the NFκb pathway appears relatively insensitive to Ca^2+^ changes even in WT HEK293 and HeLa cells.

### ROS handling is altered in TKO cells

Increased ROS levels regulate the basal activity of many transcription factors including NFAT (44,45), AP-1 (46), NFκb (47) and CREB (48). In addition, the activity of specific signaling proteins e.g. Jnk are altered by ROS (49). Therefore, we examined if ROS production is altered in the IP_3_R TKO cells. Initially, we measured the production of extracellular H_2_O_2_ using a peroxidase/amplex red assay. The results showed increased H_2_O_2_ release in both HEK293 and HeLa TKO cells under basal conditions (**Figure 4A & B**). As a positive control we treated cells with antimycin A (AA) which blocks electron flow at complex III and stimulates production of O_2_^•─^(50) that can be enzymatically converted to H_2_O_2_ by superoxide dismutase (SOD). Both WT and TKO cells showed a stimulation of H_2_O_2_ when treated with AA (Fig 4A & B). Further inhibition of the ETC with the combination of AA and rotenone blocks electron entry to the ETC and acts as a negative control. In both HEK and HeLa TKO cells the AA & rotenone cocktail was insufficient to block H_2_O_2_ release. However, in both HEK and HeLa the AA & rotenone cocktail did not eliminate the difference between WT and TKO cells suggesting an ETC independent source of H_2_O_2_ in TKO cells. We employed diphenyleneiodonium (DPI) as an isoform insensitive inhibitor of NADPH oxidase (NOX) complex assembly. In both HEK and HeLa, NOX inhibition had a more pronounced effect in TKO cells vs. WT. As predicted, inhibition of catalase activity with 3-aminotriazole (3-AT) increased H_2_O_2_ release in both HEK & HeLa TKO cells (Figure 4A&B). Inhibition of H_2_O_2_ metabolism could also contribute to the elevation of H_2_O_2_ in TKO cells. Direct measurements of the catalytic activity of glutathione peroxidase showed a small decrease that was confined to HEK293 TKO cells only (Fig S3A).

**Figure 4.**
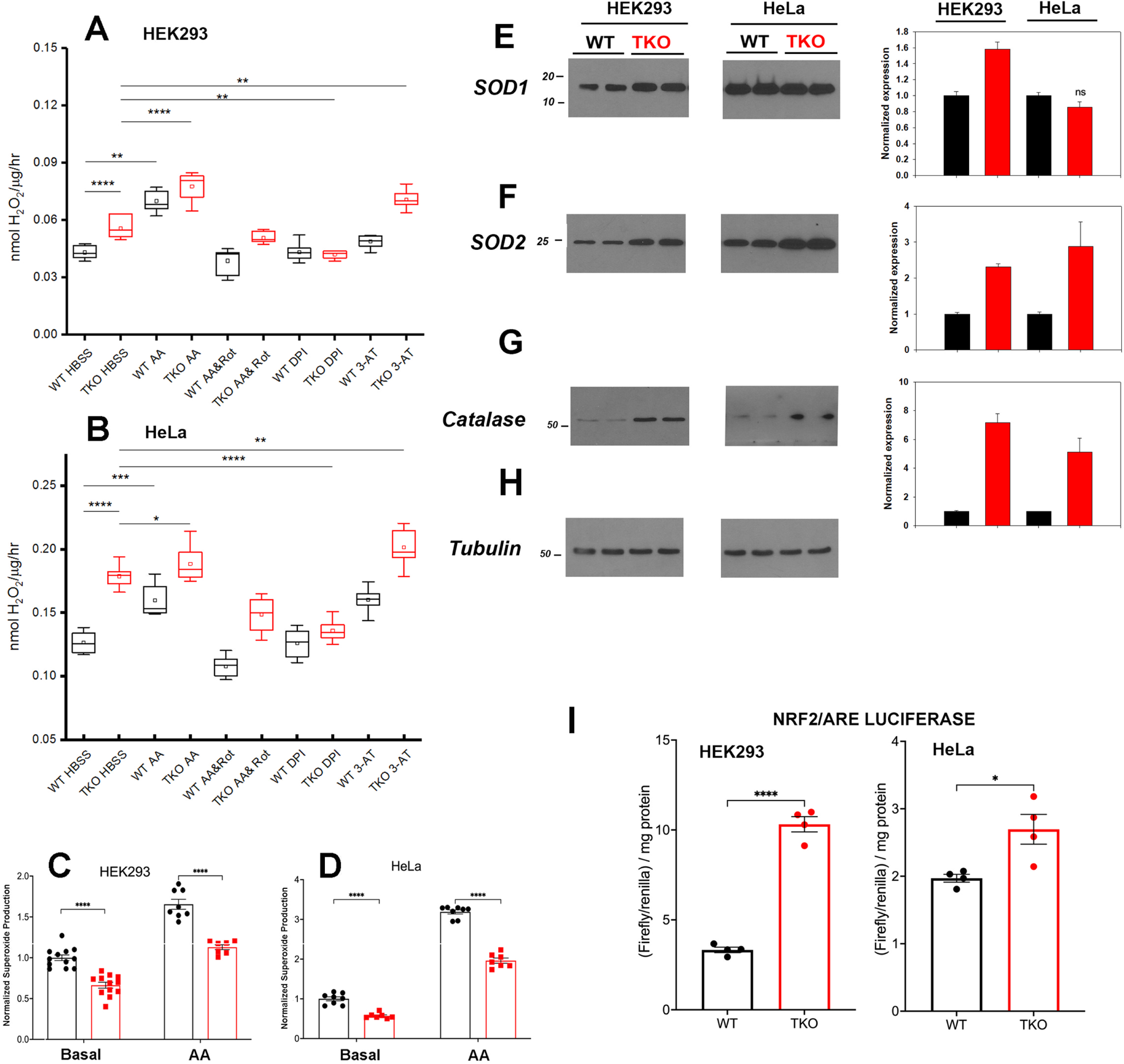
ROS handling is altered in IP_3_R TKO cells. ***Panel A*** The rate of extracellular H_2_O_2_ generation was measured with Amplex Red in WT and TKO HEK293 cells grown in 96 well plates as described in ‘Experimental Procedures’. Antimycin A (AA; 0.5μM) was used as a positive control while AA & Rotenone (Rot; 5µM), diphenyleneiodonium chloride (DPI; 5µM) and 3-aminotriazole (3-AT; 5mM) were used to inhibit the electron transport chain, NADPH oxidase and catalase respectively. The data shown are box plots showing mean (small box), median (horizontal line) 25^th^ & 75^th^ percentile (box) and 1.5 x interquartile range (whiskers). Statistics are one-way ANOVA following a normality test and subject to Tukey post-hoc analysis. *-**** represents *p* ≤ 0.05-*p* ≤ 0.0001 respectively. ***Panel B*** Conditions were as in Panel A but using HeLa WT and TKO cells. ***Panel C*** Mitochondrial superoxide was measured in WT and TKO HEK293 cells grown in 96 well plates with MitoSox. The data shown are individual measurements made in 3 separate experiments with the data normalized to the mean value of WT cells. ***Panel E-H*** Key antioxidant enzyme levels were immunoblotted in HEK293 and HeLa lysates using tubulin as the loading control. Data were quantitated and normalized to the WT unstimulated values. The data are the mean ± S.E.M. of 3 independent experiments. All data are statistically significant from WT unstimulated values at p<0.05 with only the data not significantly different marked “ns”. ***Panel E*** Superoxide dismutase-1 (SOD1). ***Panel F*** Superoxide dismutase-2. ***Panel G*** Catalase. ***Panel H*** Tubulin. ***Panel I*** HEK293 and HeLa WT (black) and TKO (red) cell lines were co-transfected with Nrf2/ARE-luciferase and *Renilla* luciferase reporter vectors. After 24h, cells lysates were assayed for promoter activity.

We attempted to quantitate superoxide (O_2_^•─^) generation directly using the mitochondrial targeted probe MitoSox (51). These results showed that O_2_^•─^ levels are decreased under basal and AA-treated conditions in both HEK293 and HeLa TKO cells (Figure 4C & D). The result would be consistent with an increased conversion of O_2_^•─^ to H_2_O_2_ by SOD in TKO cells. Indeed, levels of the cytosolic (SOD1) and the mitochondrial (SOD2) isoforms were increased in HEK293 TKO cells (Fig 4E & F). However, only the mitochondrial SOD2 isoform was elevated in HeLa TKO cells (Fig 4E & F). Catalase protein levels were also elevated in both cell types (Fig 4G). The presence of altered handling of ROS in TKO cells was further confirmed by using two additional approaches. The first was to monitor the redox status of glutathione in the cytosol and mitochondrial compartments using the genetically targeted Grx1-roGFP2 probe (52). The results showed the cytosolic glutathione pool was more reduced in the TKO cells but the mitochondrial pool was not affected (Fig S3B). The second approach was to use HyPer7-an ultra-sensitive, pH independent, H_2_O_2_ probe targeted to the cytosol (53). Using additions of H_2_O_2_ and DTT to provide maximum and minimum values, the HyPer7 probe confirmed that HEK293 TKO cells maintained a higher baseline production of H_2_O_2_ (Fig S3C) and that AA was able to further increase the signal in both WT and TKO cells. Some of the functional effects of ROS on gene transcription are mediated by the transcription factor Nrf2 which translocates to the nucleus and binds antioxidant response elements (ARE) in the promoters of genes coding for antioxidant enzymes (including catalase and SODs) (54). Measurement of luciferase promoter activity indicates that basal activity of Nrf2/ARE was increased in both HEK293 and HeLa TKO cells, consistent with enhanced ROS generation (Fig 4I).

### RNAseq analysis of HEK293 and HeLa TKO cell model

To gain a broader understanding of global baseline transcriptional changes resulting from knockout of IP3 receptors, we carried out RNAseq analysis on unstimulated WT and TKO cell models in both HEK293 and HeLa lines. For HEK293 cells a total of 58,825 gene transcripts were measured and 828 genes were identified as being differentially expressed (DEG) by edgeR analysis using a criteria of |log_2_ fold change|>1 and p value<0.05. These genes are depicted on a volcano plot in **Figure 5A** with the top 30 genes having highest statistical significance (false discovery rate p value<0.01) being colored and labeled. In order to determine if there were changes in groups of genes with related biological function, we also compared the DEGs to online curated gene sets encoding Ca^2+^ signaling components, transcription factors or mitochondrial proteins (Fig 5B). Six cell surface receptors coupled to phospholipase-C activation are in the Ca^2+^ signaling list but the biological relevance of these changes are likely to be complicated since some are upregulated (e.g. HTR2C,CCKBR) and others down regulated (e.g. EGFR,P2RY11). Six cation channels are also in the list including several members of the TRP family. This includes TRPC6, which can be directly stimulated by diacylglycerol, and is known to be involved in NFAT activation in the heart (55). All these channels, with one exception (KCNN3), were upregulated in the TKO cells. In the case of transcription factors, the list included 8 immediate early genes (underlined), all of which were downregulated under basal conditions. Only a few genes encoding mitochondrial proteins were altered. In many cases, such as acyl CoA synthetases (ACSL6, ACSL1) or respiratory complexes (NDUFA6,SCO2) the changes were in opposite directions. Transcripts for MICU2 were also decreased in TKO cells but previous studies have shown that the levels of protein were not altered (56). The over representation of DEGs in specific KEGG pathways was assessed in Fig 5C. The top 20 hits included several signaling pathways, such as phospholipase D, MAPK, and VEGF. Of particular interest, the NFκB signaling pathway was highlighted as being significantly enriched in the TKO cells which is in line with results from the NFκb luciferase reporter assays. Focal adhesion was also a highlighted KEGG pathway. Although not specifically investigated in this study, others have reported decreased motility of IP_3_R TKO HEK293 cells (6).

**Figure 5.**
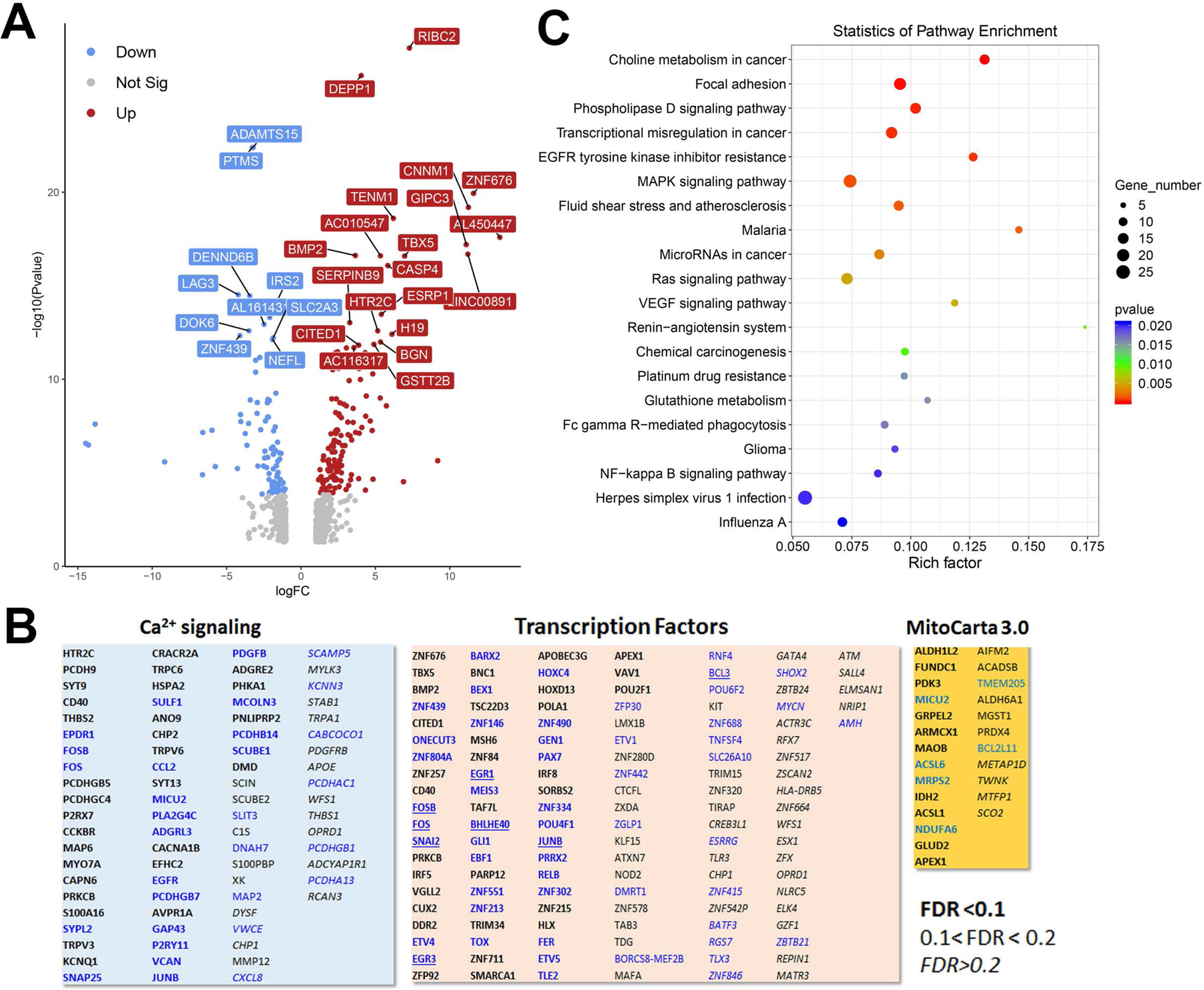
Analysis of the HEK293 RNAseq data set for Wild-type versus IP_3_R TKO cells. ***Panel A*.** Distribution of the 828 differentially expressed genes is shown as a volcano plot. The default cut off for FDR was set at 1% and |log_2_ FC| >1. The top 30 genes are indicated*. **Panel B.*** Bubble plot of KEGG enrichment analysis of DEGs. Top 20 KEGG pathways (*p* ≤ 0.05) are presented. *Y*-axis represents pathways; *X*-axis represents rich factor; (rich factor equals the ratio between the DEGs and all annotated genes enriched in the pathway). The color and size of each bubble represent enrichment significance and the number of DEGs enriched in a pathway, respectively. ***Panel C***. Manually curated gene sets for Ca^2+^ signaling pathways, transcription factors and mitochondrial genes (MitoCarta 3.0) were used to identify hits within the DEG gene list. The significance level of the gene changes measured by FDR is indicated in the inset. Genes in black were upregulated and genes in blue were down-regulated. The origin of the curated gene sets are given in ‘Materials and Methods’.

To look for cell type dependent and independent changes, we repeated the RNAseq analysis in HeLa WT and TKO cell models (**Fig 6**). For HeLa cells a total of 60,613 gene transcripts were measured and 310 genes were identified as being differentially expressed (DEG) by edgeR analysis using the same threshold criteria as used for HEK293 cells. Based on this the number of DEGs are significantly lower in HeLa cells (37.4%). These genes are depicted on a volcano plot in Fig 6A with the tope 30 genes being identified as in Fig 5A. A feature that is apparent is that many more genes are down-regulated than up regulated in HeLa cells, whereas the opposite is true for HEK293 cells. The distribution of the HeLa DEGs within the curated gene sets of Ca^2+^ signaling, transcription factors or mitochondrial proteins are shown in Fig 6B. As expected, a lower number of genes are found in each category than seen for HEK293 cells. The over-representation analysis also showed no overlap in the two cell types with a lower number of HeLa DEGs in each KEGG gene set (Fig 6C). At the single gene level with the significance criteria used, there were only 18 genes changed in common between the HEK293 and HeLa data sets (**Fig 7**). To avoid focusing on individual genes, or DEG lists based on thresholds, we utilized gene set enrichment analysis (GSEA) which uses all the genes in the RNAseq data set to look for concerted changes in biological pathways. This analysis, using the Hallmark gene set from the Molecular Signatures data base, is shown for HEK293 (Fig4.7C) and HeLa cells (Fig4.7D). More pathways were enriched than de-enriched in HEK293 cells than in HeLa cells, in line with our previous analysis of individual genes. Ten common pathways were altered in the same direction in both cell types under basal conditions. This included KRAS signaling, TNF-alpha signaling via NF-Kb, Notch signaling, hypoxia and the p53 pathway. However, there were also changes in pathways that were unique or changed in opposite directions in both cell types.

**Figure 6.**
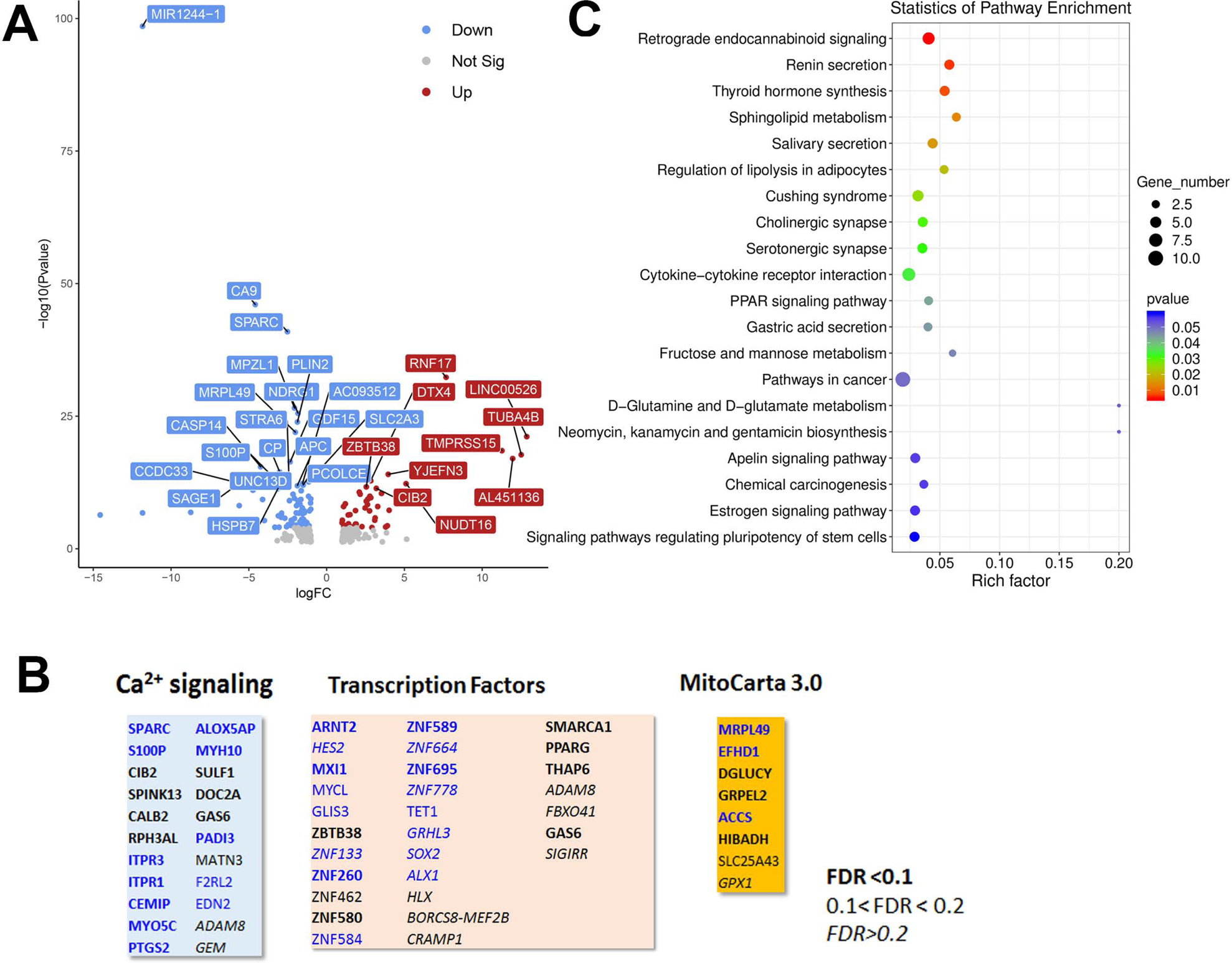
Analysis of the HeLa RNAseq data set for Wild-type versus IP_3_R TKO cells. ***Panel A*.** Distribution of the 310 differentially expressed genes is shown as a volcano plot. The default cut off for FDR was set at 1% and |log_2_ FC| >1. The top 30 genes are indicated*. **Panel B.*** Bubble plot of KEGG enrichment analysis of DEGs. Top 20 KEGG pathways (*p* ≤ 0.05) are presented. *Y*-axis represents pathways; *X*-axis represents rich factor; (rich factor equals the ratio between the DEGs and all annotated genes enriched in the pathway). The color and size of each bubble represent enrichment significance and the number of DEGs enriched in a pathway, respectively. ***Panel C***. Manually curated gene sets for Ca^2+^ signaling pathways, transcription factors and mitochondrial genes (MitoCarta 3.0) were used to identify hits within the DEG gene list. The significance level of the gene changes measured by FDR is indicated in the inset. Genes in black were upregulated and genes in blue were down-regulated. The origin of the curated gene sets are given in ‘Materials and Methods’.

**Figure 7.**
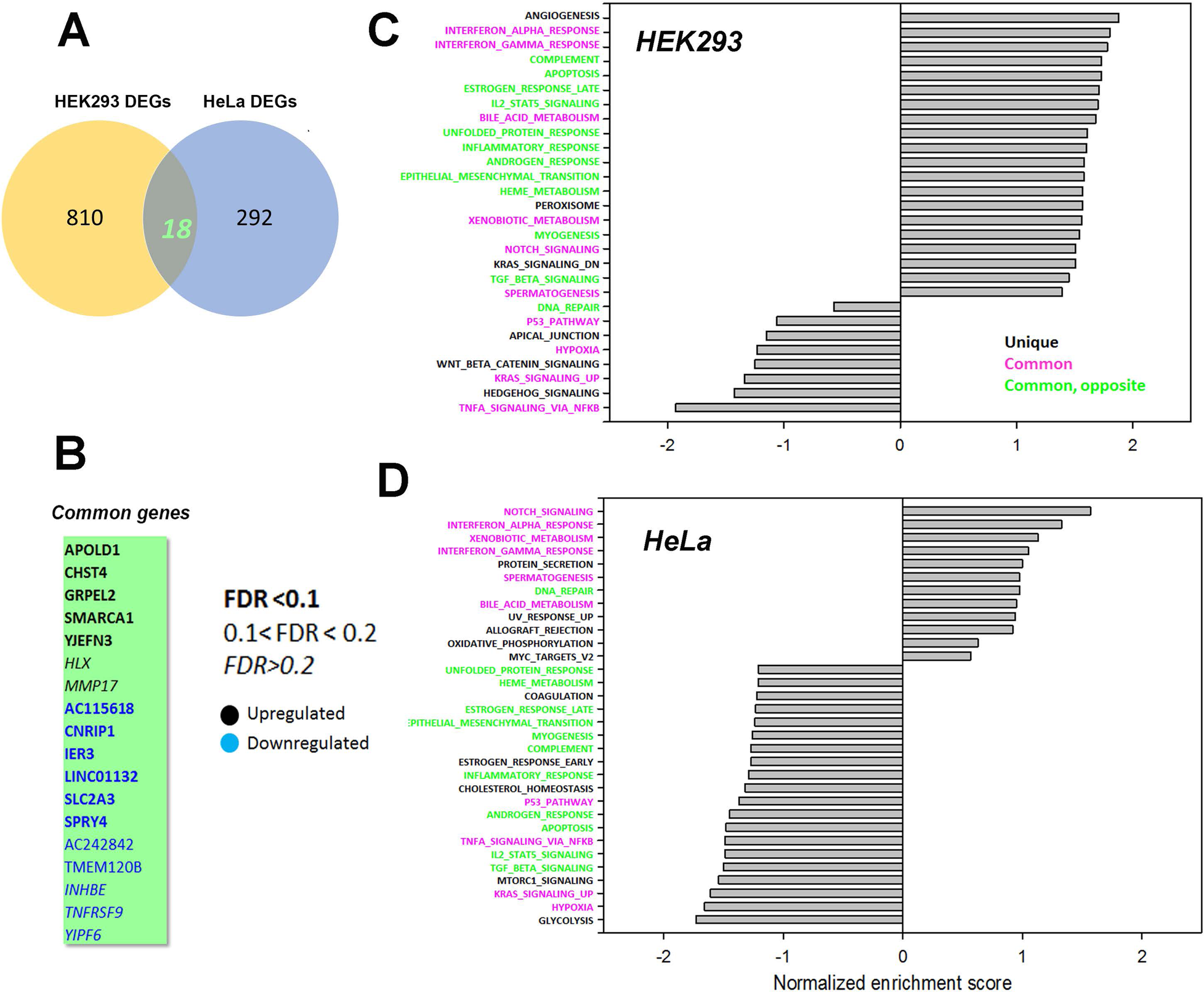
Common genes or pathways altered in the HEK293 or HeLa data sets. ***Panel A*** Venn diagram showing the common genes between the HEK293 and HeLa data sets with ***Panel B*** listing the 18 common genes. The significance level of the gene changes measured by FDR is indicated in the inset. Genes in black were upregulated and genes in blue were down-regulated. ***Panel C & D*** Gene set enrichment analysis (GSEA) was carried out using a Hallmark gene set found in the Molecular Signatures Database (http://www.gsea-msigdb.org/gsea/msigdb/collections.jsp). The statistical criteria for significance was a nominal p val < 0.05 and a false discovery rate q-val < 0.25.The top 20 enriched or de-enriched categories in the TKO cells based on normalized enrichment score (NES) are shown. Pathways in both data sets that were unique (black), common (red), or common but changing in opposite directions (green) were colored as indicated.

### Target genes of calcium-regulated transcription factors

Our focus in this study was on transcriptional changes that may result from the loss of Ca^2+^ signaling in the IP_3_R TKO cells. In particular, we have concentrated on the known Ca^2+^ dependent transcription factors e.g. NFAT, CREB, AP-1 and NFκb. Therefore, we wanted to know the contribution of target genes of these transcription factors to the RNAseq data sets. To do this we employed a publicly available transcription factor target gene data base (Harmonizome; (57)). Of the 828 DEGs in HEK293 cells, 135 genes (16.3%) were NFAT2 targets, 26 (3.1%) were CREB targets, 587 (70.1%) were AP-1 targets and 96 (11.6%) were NF-kB targets. For the 310 DEGs in HeLa cells we identified 47 genes (15.2%) as NFAT2 targets, 6 genes (1.9%) as CREB targets, 207 (66.7%) as AP-1 targets and 33 genes (10.6%) as NF-kB targets. Thus, AP-1 and NFAT were the most affected of the three transcription factors measured in TKO cells. Despite the differences in overall number of DEGs between HEK293 and HeLa cells, the percentage of altered NFAT, CREB, AP-1 and NFκB target genes were similar in both cell lines. It is important to emphasize that the RNAseq data set were obtained from unstimulated cells. NFAT and AP-1 family members can bind cooperatively to adjacent promoter sites and PKC is a known activator of AP-1 (36). This raises the possibility that, although the absence of Ca^2+^ signaling in IP_3_R TKO cells may impair the activation of NFAT, the higher basal activity of NFAT, together with the activation of Ca^2+^ -independent PKC isoforms may be sufficient to allow agonists to turn on NFAT target genes. The redundancy of transcription factor activation makes it difficult to test this hypothesis with target genes that are exclusively NFAT responsive. RT-qPCR measurements were made of two representative NFAT target genes (COX2 (58), IRS2 (59)) and ETS1, which is a transcription factor that interacts with NFAT (60,61). None of the selected genes showed agonist-mediated increases over a 6h period in HEK293 or HeLa cells (Fig S4). PMA stimulated the expression of COX2 and ETS1 in both WT HEK293 and HeLa cells (FigS4 A-D). The PMA response of COX2 was further increased in TKO cells in both cell lines, but for ETS1 was increased only in HeLa TKO cells. Baseline IRS2 mRNA levels were strongly suppressed in HEK293 TKO cells and to a lesser extent in HeLa TKO cells. This confirms findings from RNAseq data (Fig 5). Previous microarray studies on clones of HEK293 cells with high and low store-operated Ca^2+^ entry (SOCE) identified IRS2 as a putative regulator of SOCE (62), but a mechanistic and functional link between loss of IP_3_R-mediated Ca^2+^ fluxes and IRS2 remains to be established.

## DISCUSSION

Calcium signaling regulates many critical biological processes. This includes the activity of multiple transcription factors, some of which have been shown to be tuned to specific frequencies of Ca^2+^ oscillations (63,64). In the present study we have examined the behavior of key Ca^2+^-dependent transcription factors in the setting of a complete loss of IP_3_-linked Ca^2+^ signaling using two human cancer cell lines that have been genetically deleted of all three IP_3_R isoforms. The main findings of this study are that TKO cells show: a) an elevated baseline activity of several transcription factors (e.g. NFAT, CREB, AP-1); b) a loss of agonist-mediated activation of NFAT, but not of other transcription factors (e.g. CREB); c) an increased reliance on the Ca^2+^-insensitive forms of PKC for downstream signaling and d) an altered handling of ROS.

Our studies confirm earlier reports that TKO cells have defective agonist-mediated NFAT activation in chicken DT40 cells (2), mouse B-lymphocytes (4) and mouse embryonic stem cells (8). The two new findings made in the present study are an elevated baseline NFAT activity in both HEK293 and HeLa TKO cells, and an enhanced sensitivity of NFAT to pharmacological elevation of cytosolic Ca^2+^, observed in HEK293 but not in HeLa TKO cells. There is no evidence to suggest that baseline levels of cytosolic Ca^2+^ or ionophore-mediated Ca^2+^ release are any different in TKO cells (6,12). Therefore, it is more likely that the Ca^2+^ sensitivity of the CaN/NFAT signaling pathway to baseline and elevated Ca^2+^ is enhanced in HEK293 TKO cells. A possible molecular mechanism may involve decreased levels of RCAN1, an endogenous inhibitor of CaN (Fig 1D). RCAN1 levels and ionophore sensitivity were unchanged in HeLa TKO cells, suggesting that the mechanism of baseline elevation of NFAT activity in HeLa cells is likely to be different from HEK293 cells and may be related to the different complement of NFAT isoforms in the two cell lines (23).

We also found increased baseline activity of CREB in both HEK293 and HeLa TKO cells, but again only the HEK293 TKO cells showed an enhanced response to ionophore that was suppressed by CsA (Fig 2A). Like NFAT, CREB activation is also linked to CaN, independently of ser133 phosphorylation, since the nuclear translocation of the CREB co-activator CRTC1-3 requires dephosphorylation by CaN (30). The upstream kinase regulation of CRTCs is different in HeLa cells (31) and therefore the different sensitivity of CREB to ionophore in the two TKO cell lines is not unexpected. Agonist-induced increases of CREB ser-133 phosphorylation is maintained in both TKO cell lines, and our studies with PKC inhibitors and siRNA suggests that this is mediated in part by the Ca^2+^ insensitive PKCδ isoform. An increased reliance on Ca^2+^-insensitive PKC isoforms may be an important adaptive mechanism maintaining down-stream signaling in the absence of Ca^2+^ changes. In WT cells with a Ca^2+^ signal, competition for membrane-bound DAG favors the translocation of classical rather than the novel PKC isoforms (65). When the Ca^2+^ signals are eliminated with BAPTA loading, only the novel PKC isoforms show translocation (65). Thus, the increased reliance on the novel PKC isoforms in TKO cells may reflect the outcome of this competition without necessarily involving altered DAG levels or additional regulatory mechanisms.

Another transcription factor that showed increased baseline promoter activity in TKO cells was AP-1 and this was associated with elevated protein levels of some members of the fos and jun family members (e.g. c-jun in HEK293 cells). The rapid induction of c-fos by elevated Ca^2+^ is a well documented effect in many cell types (18), but in the human cancer cell lines studied here, c-fos was already constitutively expressed under basal conditions and showed only small increases with agonist in WT and TKO cells. However, in WT HEK293 cells fosB was absent under baseline conditions and was markedly induced by agonist only in WT and not TKO cells. This suggests that fosB induction may also require NFAT activation. Studies in cultured monocytes have shown that induction of fosB is dependent on NFAT4 activation (66). Since dimeric members of the fos family have no transcriptional activity by themselves, they must bind with other partners, which include NFATs (66). Further studies are required to determine the transcriptional consequences of the loss of fosB induction in HEK293 TKO cells.

Increased production of ROS may contribute to the baseline elevation of many of the transcription factors examined in the present study (44–49). Our main observations on ROS were that H_2_O_2_ production was increased and superoxide production was decreased in both TKO cell lines, consistent with enhanced dismutation by increased SOD enzyme concentrations (Fig 4E&F). It is important to stress that the TKO cells are not undergoing ‘oxidative stress’, since the cytosolic glutathione pool was actually more reduced than WT cells (Fig S3B). The changes appear unrelated to the total NADP^+^/NADPH ratio since this was not significantly changed in TKO cells (12). Our data suggest that redox balance is maintained, in part, because the increased production of ROS is accompanied by increased expression of antioxidant enzymes, including catalase and superoxide dismutase, suggesting that increasing H_2_O_2_ for signaling purposes may be an important adaptation in the absence of ER Ca^2+^ signaling. The transcription of these enzymes are regulated by the redox-sensitive transcription factor Nrf2 (54), which also shows increased activity in TKO cells (Fig4I). Inhibitor studies point to the possible involvement of an NADPH oxidase isoform, but further work is required to establish the exact source(s) of increased ROS in TKO cells. A previous study in DT40 TKO cells also concluded that ROS production was increased and that the total glutathione pool was more reduced, but the results differed in that SOD2 was decreased and catalase was unchanged (10).

A predicted consequence to maintaining a higher basal transcription factor activity would be an increased expression of the corresponding target genes. Using NFAT2 as an example 54% and 49% of the target genes were significantly upregulated in HEK293 and HeLa TKO cells respectively. It is unlikely that there is a simple relationship between increased baseline activity of a few Ca^2+^ dependent transcription factors and the overall DEGs, since each gene can be activated by multiple transcriptional activators/repressors. The functional consequences of maintaining a higher basal activity of transcription factors for agonist-mediated responses are unclear. Presumably, in wild-type cells the concerted effects of the Ca^2+^ and DAG signals allow the cells to transition from a resting to a stimulated state (**Figure 8**). We hypothesize that the maintenance of a higher basal activity in TKO cells may be a mechanism to allow signaling pathways to be fully activated by the DAG signal alone. Further work is necessary to validate this hypothesis.

**Figure 8.**
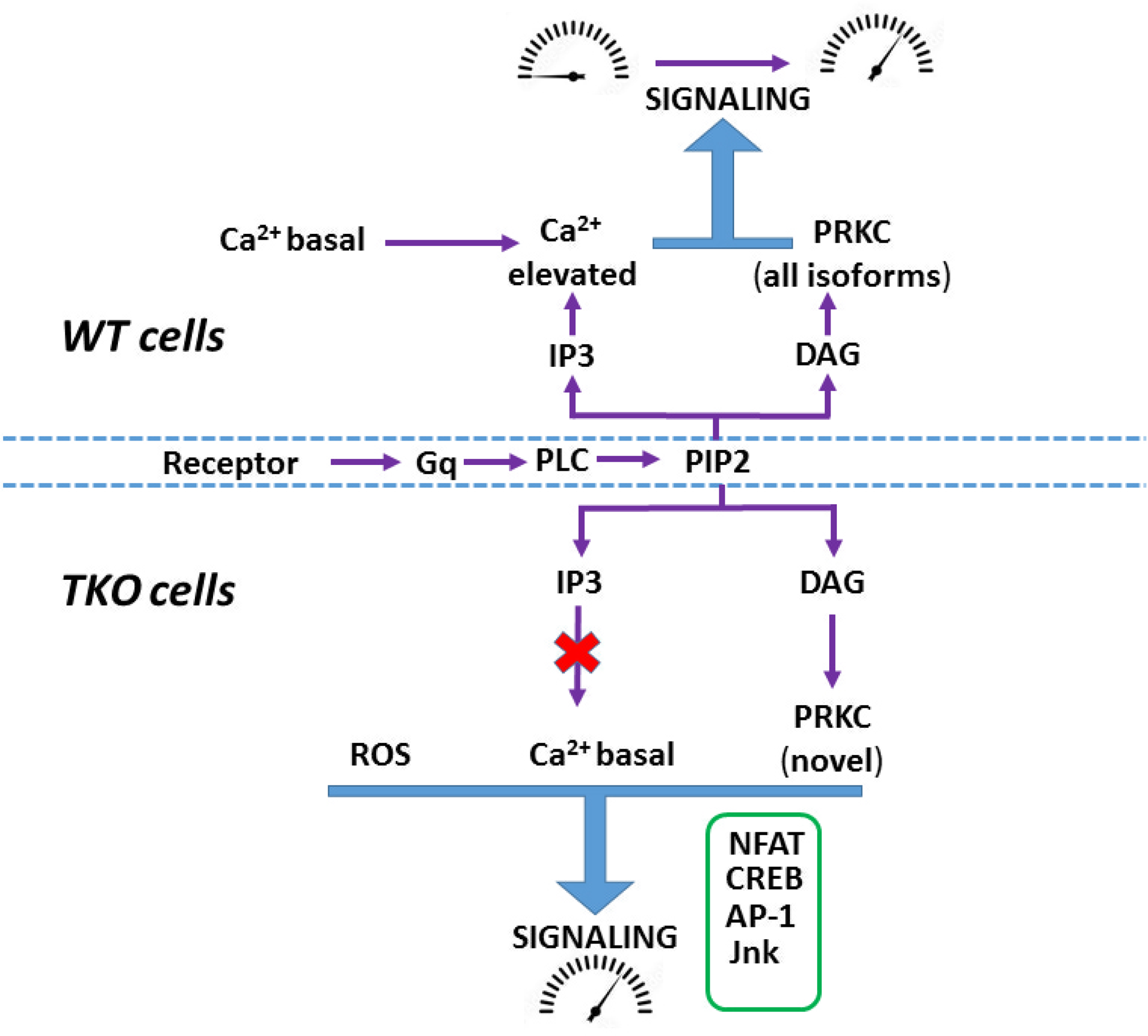
Hypothetical scheme showing agonist-mediated activation of signaling in WT and TKO cells. In WT cells both Ca^2+^ mobilization and DAG formation mediate the transition of signaling pathways from ‘resting’ to an ‘activated’ state. In TKO cells many signaling pathways have adapted by: a) adjusting their Ca^2+^ sensitivity to be active at resting Ca^2+^ (e.g. NFAT);b) Increasing their reliance on the DAG signal alone (e.g. CREB via PKCδ) and c) becoming sensitized to increased ROS (e.g. Jnk, AP-1).

Our studies challenge the widely held belief that Ca^2+^ signaling is essential for cell function and suggest that this may not be valid under all conditions. Deletions in IP_3_Rs are lethal when this occurs in tissues where only one IP_3_R isoform predominates (e.g. IP_3_R1 in brain), or in organisms expressing only one isoform (e.g. Drosophila) (67). However, where all 3 IP_3_R isoforms are present initially and are sequentially deleted, the conditions may be more favorable for the development of adaptive mechanisms to tolerate the loss of Ca^2+^ signaling. The ability of TKO human cancer cell lines to grow and divide in the absence of any Ca^2+^ signaling can probably be attributed to the maintained activity/redundancy of important signaling pathways (e.g. CREB, MAPK etc) required for growth. However, the finding that some TKO cells grow more slowly is an indication that there are Ca^2+^ regulatory step(s) that cannot be completely circumvented by adaptive mechanisms. Understanding the molecular basis of these adaptive mechanisms may provide valuable insights into how cells bypass the need for Ca^2+^ signaling in such fundamental processes such as cell division, or in genetic diseases linked to IP_3_R dysfunction.

## EXPERIMENTAL PROCEDURES

### Cell lines and culture

All cells were cultured at 37°C and 5% CO_2_ in Dulbecco’s modified Eagle’s medium (DMEM) supplemented with 5% fetal bovine serum, 1% penicillin/streptomycin and 0.25µg/ml amphotericin B. HEK293 IP_3_R TKO and HeLa IP_3_R TKO cells were the kind gift of Drs. David Yule (5) and Katsuhiko Mikoshiba (7), respectively.

### Luciferase reporter assays

2 x 10^5^ cells were plated in 12-well dishes and grown to 70-80% confluency. The reporter constructs used were as follows: pGL3-NFAT luciferase, Addgene #17870; pGL3-3xAP1, Addgene #40342; pGL4.29-CRE, Promega #E8471; pGL4.32-NFkb-RE, Promega #E8491. Cells were transfected with 1ug DNA/well of reporter gene-luciferase and 0.2ug/well of *Renilla* luciferase (a kind gift of Dr. Makarand Risbud) using Lipofectamine 3000 (ThermoFisher Scientific). After 24h, cells were subjected to various treatments for 4h and then lysed with 1X Firefly Luciferase assay buffer (Biotium Corporation). A Luciferase Assay Kit 2.0 (Biotium Corporation) was used to measure firefly and *Renilla* luciferase activity on a Synergy Neo2 (Biotek) plate reader.

### SDS-PAGE / Western blotting

To prepare lysates, cells incubated in DMEM were washed in PBS and lysed in a 0.25ml of WB buffer containing 1% Triton X-100, 50mM Tris/HCl pH 7.8, 150mM NaCl, 2mM sodium orthovanadate, 10mM sodium pyrophosphate, 20mM NaF, and a 1x dilution of a complete protease inhibitor mixture (Roche Diagnostics). The lysates were centrifuged at 12,000xg for 10min. The supernatants were denatured in SDS sample buffer. Lysates were boiled at 100°C for 5min, then stored at −20°C until use. Unless otherwise noted, 40µg of protein were run at 100V for 90min on 10% polyacrylamide gels and transferred at 100V for 60min onto nitrocellulose membranes. Polyvinylidene difluoride membranes were used in the specific case of LC3 immunoblotting. Membranes were blocked in TBST supplemented with 5% BSA for 1h at room temperature. Following 3 x 15min washes in TBST, the primary Ab was added for 16h at 4°C. All Abs were used at a dilution of 1:1000 and were obtained from Cell Signaling. Membranes were developed using ECL reagent (ThermoFisher). When necessary membranes were stripped using a buffer containing SDS (2% w/v), Tris (62.5mM;pH6.8) and β-mercaptoethanol (100mM).

### PKCδ knockdown

2 x 10^5^ cells were plated in 60mm dishes, incubated at 37°C, and grown to 70-80% confluency. Cells were transfected with either PKCδ siRNA, non-targeted negative control siRNA, or Cy3 fluorescent positive control siRNA (OriGene Technologies Inc., Rockville, MD) using a Lipofectamine RNAiMAX kit (Thermo Fisher Scientific) then incubated at 37C for 48h.

### RNAseq analysis

RNA was isolated using Direct-zol RNA Miniprep kit (Zymo Research, cat no. R2052) according to manufacturer’s instructions and yielded samples with RIN values >7. Poly-A RNA sequencing library was prepared following Illumina’s Tru-Seq-standard-mRNA sample preparation protocol. Poly(A) tail-containing mRNAs were purified using oligo-(dT) magnetic beads with two rounds of purification. After purification, poly(A) RNA was fragmented using divalent cation buffer in elevated temperature. Paired-ended sequencing was performed on Illumina’s NovaSeq 6000 sequencing system. Reads were processed to remove adapter contamination, verified for sequence quality, and then assembled and mapped to the human genome. All library construction, sequencing and data processing was performed by LC Sciences, Houston, TX. The differentially expressed mRNAs were selected with log2 (fold change) >1 or log2 (fold change) <-1 and with statistical significance (adjusted p value < 0.05) by R package edgeR (68).

### Analysis of transcriptomic data

The volcano plots were drawn using the ‘Volcano plots’ feature in the toolbox of the Galaxy web site (https://usegalaxy.eu/). Visualization of pathway enrichment of genes using the KEGG compendium of pathways was supplied by LC Sciences. Manual curation of differentially expressed genes (DEG) was done with the following data sets: calcium signaling (69)(https://www.uhlenlab.org/cagedb/), transcription factors (70)(http://humantfs.ccbr.utoronto.ca/allTFs.php) and MitoCarta3 (71) (https://personal.broadinstitute.org/scalvo/MitoCarta3.0/human.mitocarta3.0.html). Data sets of target genes for the transcription factor NFAT, CREB, AP-1 and NF-Kb were downloaded from the Harmonizome web site (https://maayanlab.cloud/Harmonizome/).Gene Set Enrichment Analysis (GSEA) (72) was performed by GSEA software (v 4.1.0) using the Hallmark gene set documented in the MSigDB-C5 (v 7.3).

### RT-qPCR

Cells were seeded at 5×10^5^ cell/well on a six-well plate and grown overnight at 37°C. After treatment, total RNA was purified using a Direct-zol RNA Microprep kit (Zymo Research, cat no. R2052) according to manufacturer’s instructions. Genomic DNA was eliminated by on-column digestion with DNaseI. RNA quality and quantity were measured using a NanoDrop. A total of 200ng RNA was reverse transcribed using 1µl reverse transciptase (200U/ml, APEXbio, cat. no. K1071). qPCR reactions were performed in triplicate using HotStart 2X SYBR qPCR Master Mix, APEXbio, cat. no. K1070) according to the manufacturer’s instructions on a QuantStudio5 (Applied Biosystems). Relative gene expression was normalized using either GAPDH or Actin as a housekeeping gene. The sequences of the primers used are given below:

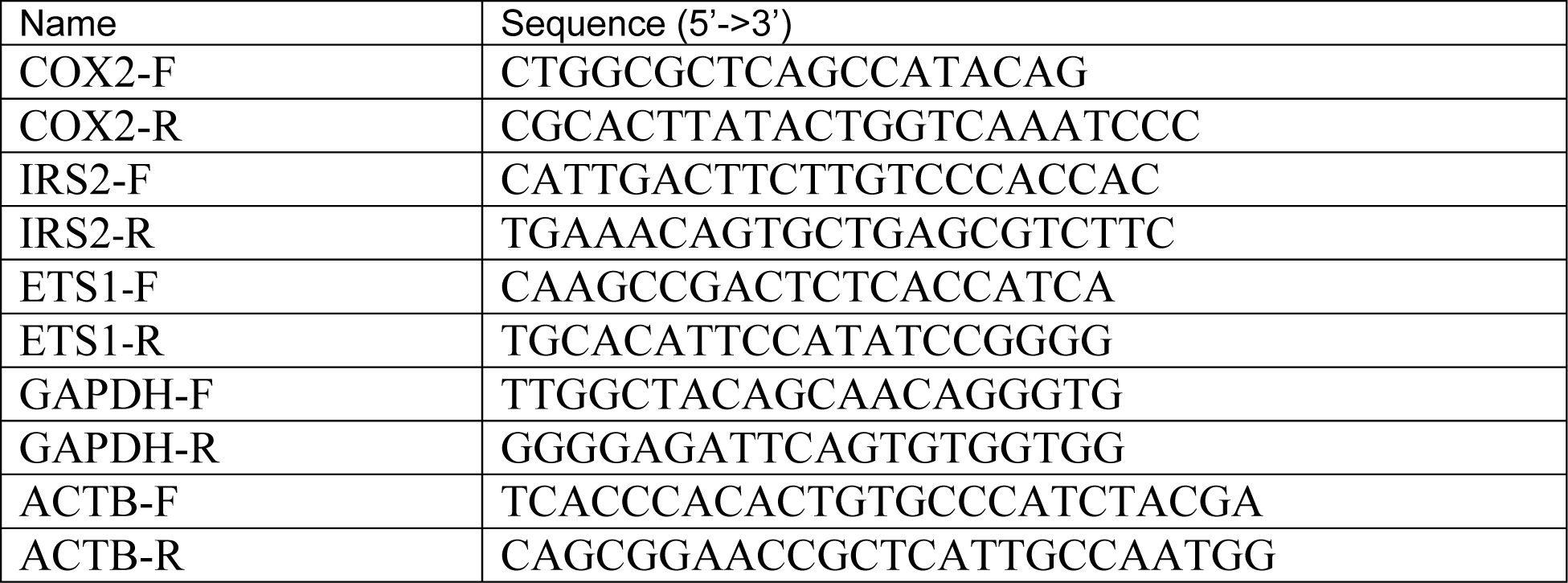

### ROS & redox measurements

Extracellular H_2_O_2_ was measured using Amplex Red reagent (ThermoFisher; A12222). 5 x 10^5^ cells/well were seeded on black, clear bottom 96-well plates and grown in DMEM overnight. Medium was aspirated from all wells and replaced with 100µl Amplex red assay buffer (HBSS, 10µM Amplex Red, 0.4U/ml horseradish peroxidase). Fluorescence was measured every 5min for 1h using a Synergy Neo2 plate reader (Biotek Instruments) set at Ex 535nm and Em 590nm. After 1h, wells were aspirated, cells were lysed using 50µl WB buffer and assayed for protein. Absolute values were estimated from a H_2_O_2_ standard curve. Mitochondrial superoxide (O_2_^•─^) was measured using a MitoSOX Red mitochondrial superoxide indicator (ThermoFisher; M36005). 5 x 10^5^ cells/well were seeded on black, clear bottom 96-well plates and grown in DMEM overnight. Medium was aspirated from all wells and replaced with 150µl MitoSOX assay buffer (HBSS, 1µM MitoSOX Red reagent, 5mM glucose, 1mM sodium pyruvate). Fluorescence was measured immediately and every 20min for 1h using a Synergy Neo2 plate reader (Biotek Instruments) set at Ex 510nm and Em 595nm. After 1h, wells were aspirated, cells were lysed using 50µl WB buffer and assayed for protein. Intracellular H_2_O_2_ was assessed using cells transiently transfected with cytosol-targeted pCS2+HyPer7-NES (Addgene #136467). Fluorescence ratios Ex490/Ex405nm and Em 535 were obtained every 30s for 30min using the Synergy Neo2 plate reader. At the end of the experiment, sequential additions of dithiothreitol (DTT) were made to establish the Ex490/405nm minimum value representing fully reduced probe. Subsequently, sequential H_2_O_2_ (200µM) washes were performed until a stable Ex490/Ex405nm maximum ratio was achieved representing a fully oxidized probe. Results are presented as (R-R_DTT_)/(R_H2O2_-R_DTT_).Glutathione peroxidase was assayed with 1ug cell lysate in 96 clear well plates using an assay buffer containing 6mM GSH, 1.25mM NADPH and 500U glutathione reductase. After mixing, 100µl 7mM H_2_O_2_ was added and the absorbance of NADPH read at 340nm using a Synergy Neo2 plate reader every 10s for 5min. Rates of NADPH oxidation/mg protein were calculated from linear regions of the graphs.

### Live cell imaging

Epifluorescence imaging of GSSG:GSH was carried in cells transiently transfected with cytosol and mitochondrial matrix-targeted Grx1roGFP2 (PMC8455442) Lipofectamine 3000 (ThermoFisher L300000) according to manufacturer’s instructions. 24-36hrs post transfection, cells were pre-incubated in HBSS supplemented with 2% BSA. For imaging, cells were washed and transferred to a similar solution containing 0.25% BSA. Imaging was carried out using a back-illuminated electron multiplying charge-coupled device (EMCCD) camera (EvolveEM 512×512 pixel. Photometrics), mounted to an Olympus IX70 inverted microscope equipped with a Sutter DG4 light source. Fluorescence excitation was achieved with a custom dichroic derived from Chroma #59022 modified to enhance short wavelength excitation and bandpass filters specific to fluorophores. Grx1roGFP2 was imaged with dual excitation of 402/15 & 485/15nm. Analysis was carried out using the FIJI variant of ImageJ (NIH, version 1.52). Images were subject to background subtraction and registration (MultiStackreg plugin, registration matrices applied equally to all wavelengths). GSSG:GSH was calculated using the formula (R-Rmin)/(Rmax-Rmin) where Rmin & Rmax values were obtained via complete reduction (DTT) and oxidation (H_2_O_2_) of Grx1roGFP at the end of each experiment. Excitation frequencies in Grx1roGFP2 are reversed, short wavelength/long wavelength to ensure that oxidation of both Grx1roGFP2 and HyPer7 causes an increase in ratio.

### Data collection and analysis

Unless otherwise noted, the data are expressed as means ± SEM of three independent experiments, each performed in technical triplicate. GraphPad Prism 8.0 (GraphPad Software, San Diego, CA, USA) was used to generate plots and to perform statistical analysis.

## ACKNOWLEDGEMENTS

This work was supported by NIH RO1 grants GM132611 (SKJ), GM151536 (GH) and 5R35GM147191 (MT). MPY and DMB were supported by an NIH NIAAA training grant T32-AA007463.

## Abbreviations

[Ca^2+^]c: cytosolic [Ca^2+^]
CaN: Calcineurin
PKC: protein kinase C
ETC: electron transport chain
COX2: cyclooxygenase-2
ETS1: E26 transformation specific sequence-1
IRS-2: insulin receptor suabstrate-2
NFAT: nuclear factor of activated T-cells
CREB: cAMP response element-binding protein
AP1: activator protein-1
NFκb: Nuclear factor kappa-light-chain-enhancer of activated B cells

---

**Supplementary Fig S1.**
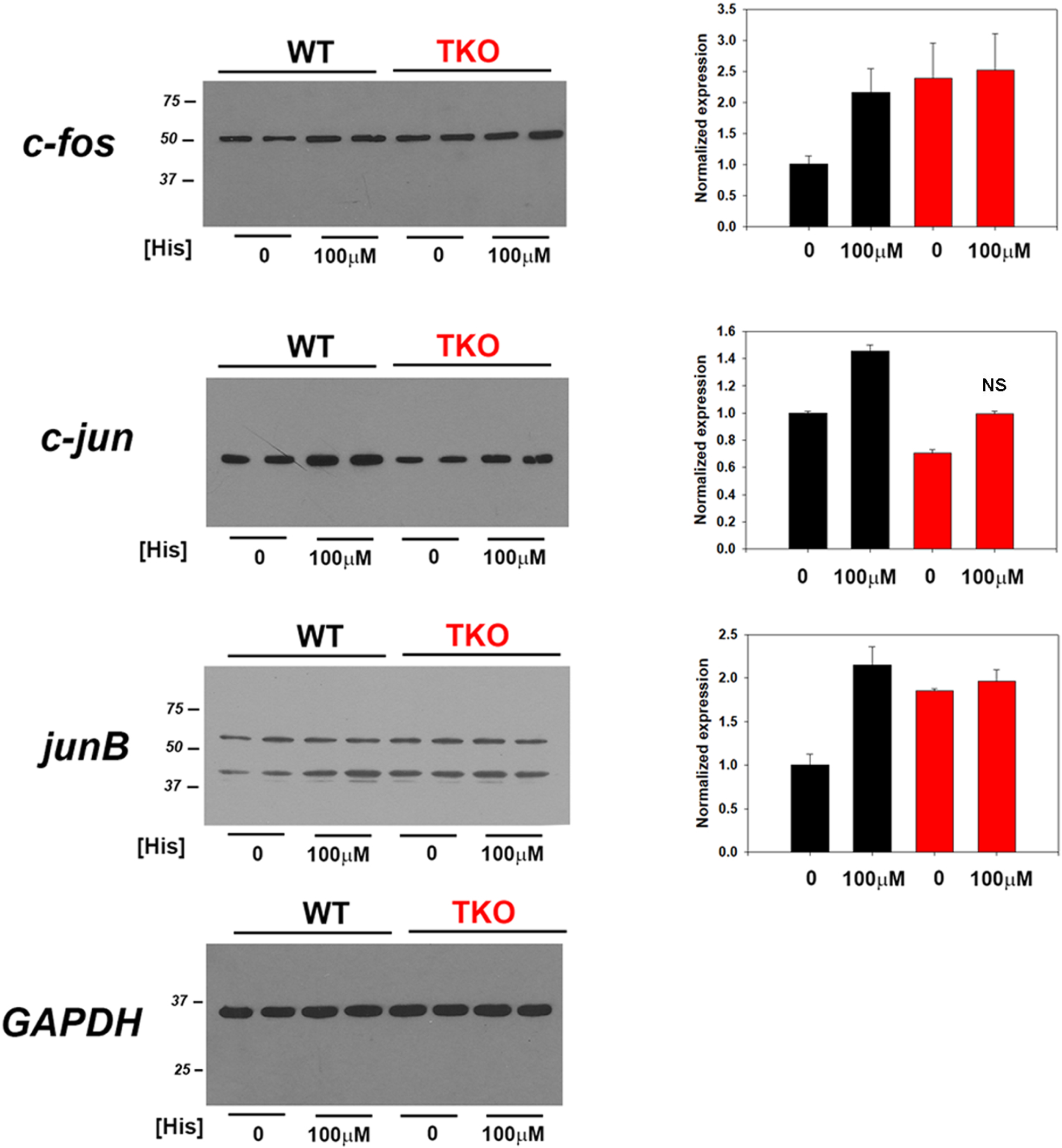
Fos and Jun protein changes in HeLa TKO cells. Lysates were prepared from HeLa WT and TKO cells treated with 100µM Histamine (His) for 4h. Samples were immunoblotted for the indicated AP-1 family members and quantitated using GAPDH as a loading control. Data were normalized to the WT unstimulated values. ***Panel A*** c-fos; ***Panel B*** c-jun; ***Panel C*** junB; ***Panel D*** GAPDH. The data shown are the mean ± S.E.M.of 3 independent experiments. All data are statistically significant from WT unstimulated values at p<0.05 with only the data not significantly different marked “ns”.

**Supplementary Fig S2.**
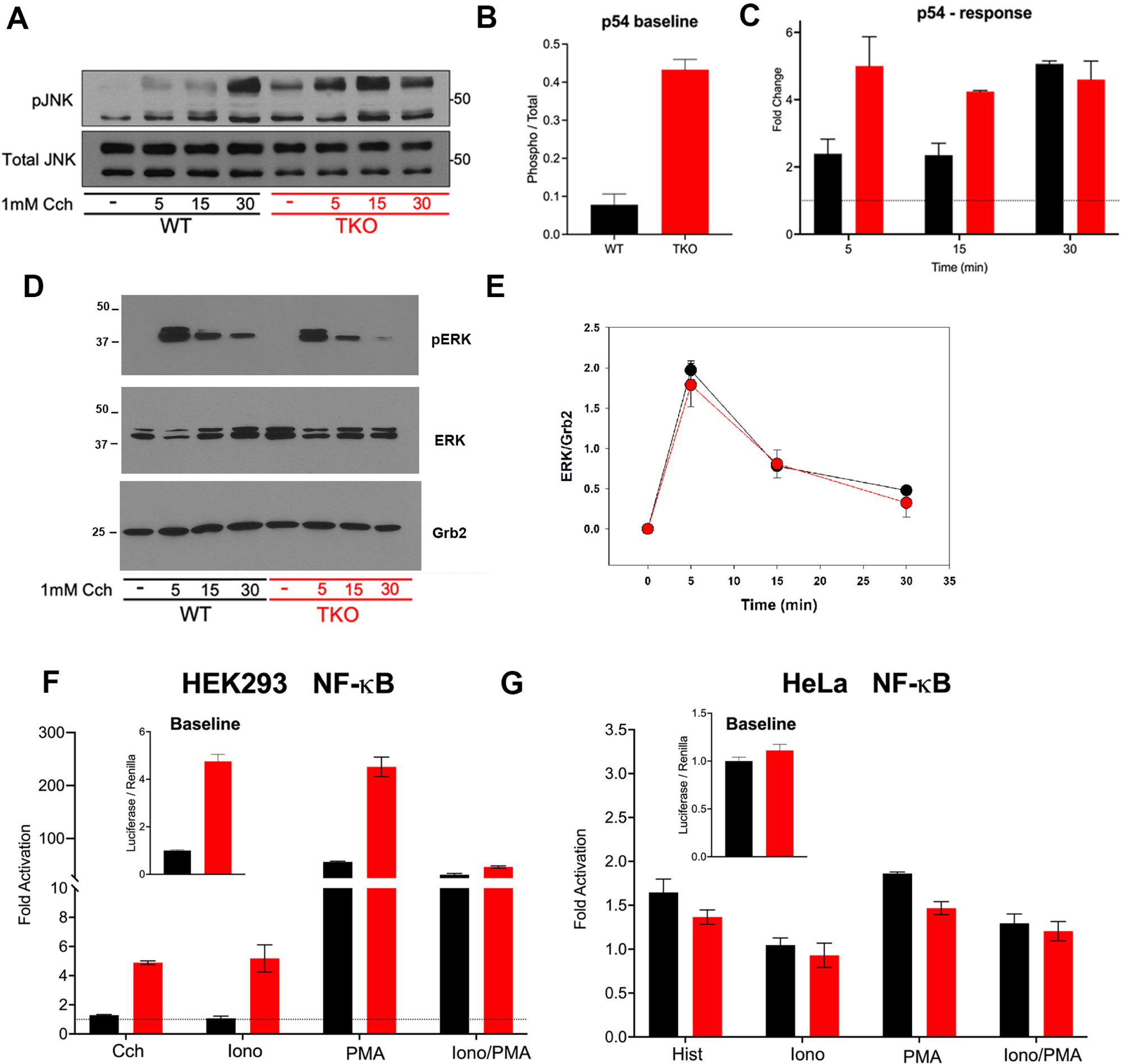
Jnk and ERK1/2 changes in HEK293 TKO cells. ***Panel A*** shows immunoblots of p-Jnk and total Jnk in WT and TKO HEK293 cells. The baseline (***Panel B***) and stimulation by 1mM carbachol (Cch) is quantitated for the p54 isoform of Jnk in 3 independent experiments (***Panel C***). ***Panel D*** shows immunoblots of p-ERK1/2 (p42/p44), total ERK and Grb2 (used as a loading control). Quantitation of the data is shown in ***Panel E***. HEK293 (***Panel F***) and HeLa (***Panel G***) were cotransfected with the NFkb luciferase reporter and Renilla luciferase constructs. Treatment conditions were as described for Fig 1.

**Supplementary Fig S3.**
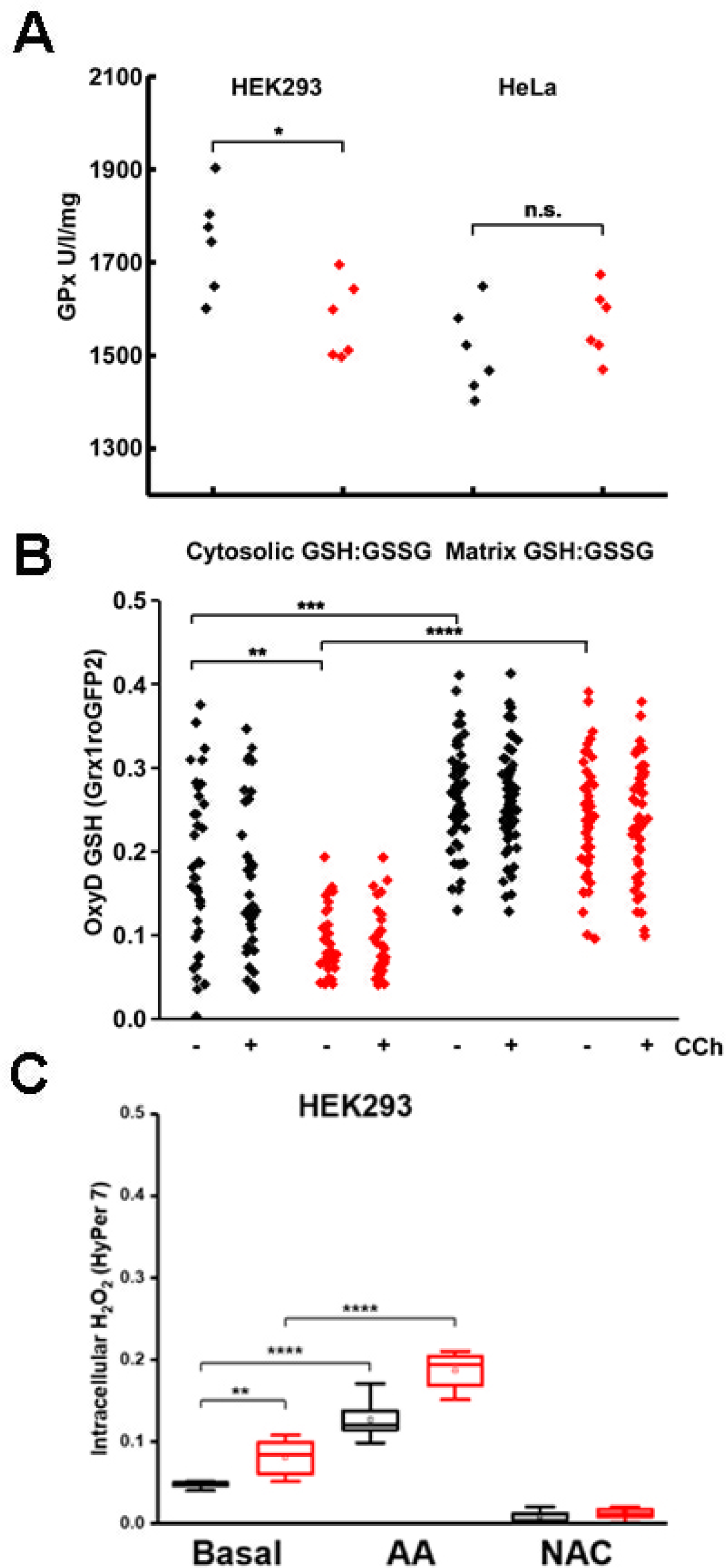
Glutathione levels and glutathione peroxidase activity in HEK293 TKO cells. ***Panel A*** shows an assay of glutathione peroxidase activity in Units per liter per mg protein (GPx U/l/mg) in lysates derived from WT (black) and TKO (red) HEK293 (left) and HeLa (right) cells as detailed in Experimental procedures. * Represents *p* ≤ 0.05, data points represent individual technical replicates. ***Panel B*** Shows the oxidized to reduced glutathione (OxyD; GSH:GSSG) measured with Grx1roGFP2 targeted to the cytosol (left) or mitochondrial matrix under basal or agonist simulated conditions (Charbachol, CCh 1mM 10 minutes). **,*** & **** represent *p* ≤ 0.05,0.01 & 0.001 respectively in a Kruskal Wallis test. ***Panel C*** The concentration of intracellular H_2_O_2_ was measured with HyPer7 targeted to the cytosol transiently transfected into HEK293 WT and TKO cells growing in 96well plates as described in “Experimental Procedures”. Cells were treated with Antimycin A (AA; 0.5μM), and N-acetyl cysteine (NAC; 1mM) as positive and negative controls respectively. Data presented as (R-Rmin)/(Rmax-Rmin). Statistics as in Fig 4A&B.

**Supplementary Fig S4.**
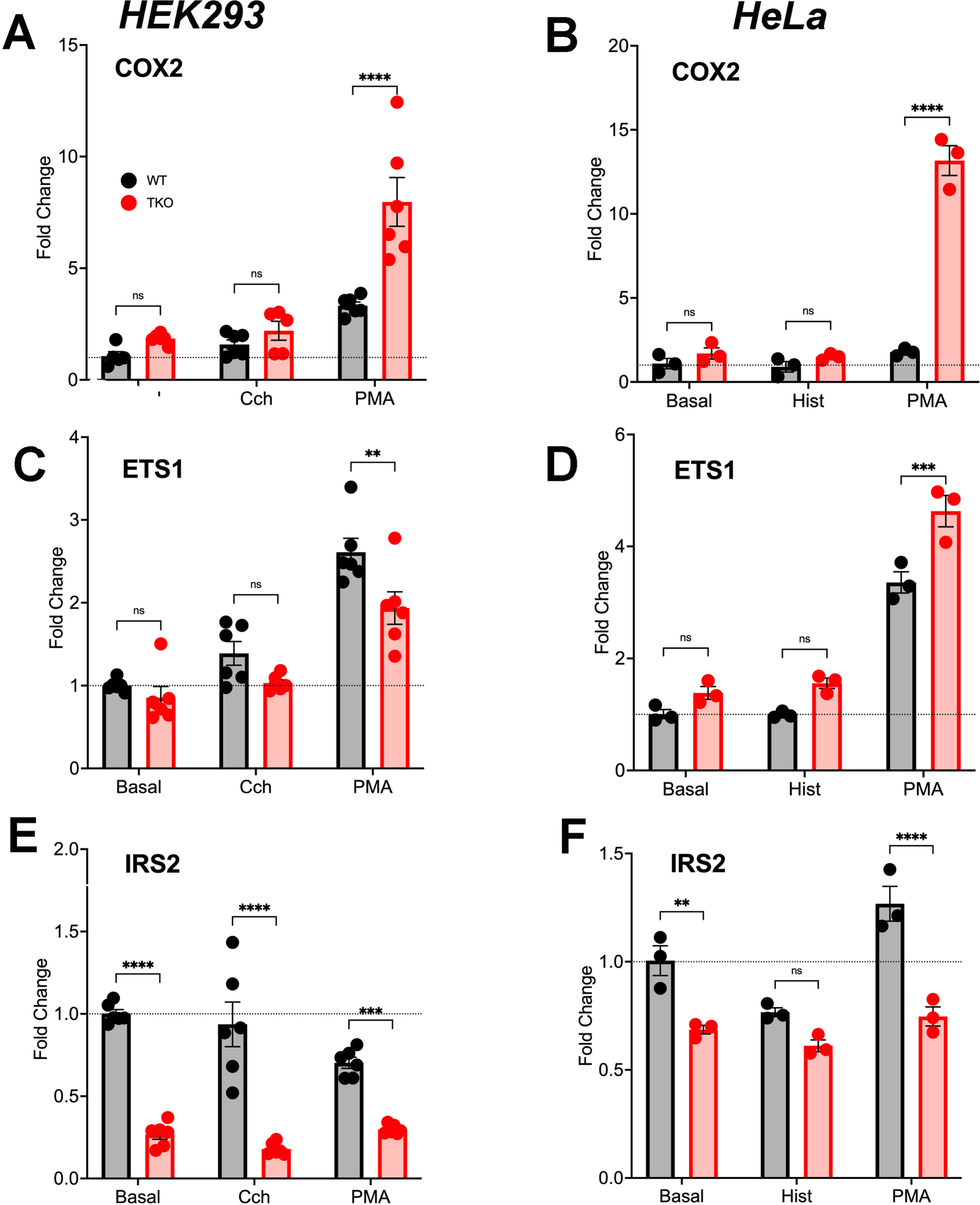
RT-qPCR and target genes of transcription factors in HEK293 and HeLa TKO cells. RT-qPCR was performed using the primers for COX2, ETS1 and IRS2 genes as described in ‘Experimental Procedures’. The experiments were done on HEK293 (Panels A,C&E) or HeLa (Panels B,D & F) stimulated for 6h with agonist (carbachol (Cch) 1mM; histamine (His) 100uM) or PMA (100nM). Relative expression levels of target genes were analyzed by the 2 ^−ΔΔCt^ method using either actin or GAPDH as a housekeeping gene. Data are expressed as fold-change compared to the untreated WT control. Each experiment was performed twice using three technical replicates. Data was expressed as the mean ± SD and significance between WT and TKO cells was assessed by Student’s t-test.

## REFERENCES

1. Berridge, M. J., Bootman, M. D., and Roderick, H. L. (2003) Calcium: Calcium signalling: dynamics, homeostasis and remodelling. Nature review Molecular Cell Biology 4, 517–529

2. Sugawara, H., Kurosaki, M., Takata, M., and Kurosaki, T. (1997) Genetic evidence for involvement of type 1, type 2 and type 3 inositol 1,4,5-trisphosphate receptors in signal transduction through the B-cell antigen receptor. EMBO J. 16, 3078–3088

3. Ouyang, K., Leandro Gomez-Amaro, R., Stachura, D. L., Tang, H., Peng, X., Fang, X., Traver, D., Evans, S. M., and Chen, J. (2014) Loss of IP3R-dependent Ca2+ signalling in thymocytes leads to aberrant development and acute lymphoblastic leukemia. Nature communications 5, 10.1038/ncomms5814

4. Tang, H., Wang, H., Lin, Q., Fan, F., Zhang, F., Peng, X., Fang, X., Liu, J., and Ouyang, K. (2017) Loss of IP3 Receptor-Mediated Ca(2+) Release in Mouse B Cells Results in Abnormal B Cell Development and Function. J Immunol 199, 570–580

5. Alzayady, K. J., Wang, L., Chandrasekhar, R., Wagner, L. E., 2nd, Van Petegem, F., and Yule, D. I. (2016) Defining the stoichiometry of inositol 1,4,5-trisphosphate binding required to initiate Ca2+ release. Science signaling 9, 10.1126/scisignal.aad6281

6. Yue, L., Wang, L., Du, Y., Zhang, W., Hamada, K., Matsumoto, Y., Jin, X., Zhou, Y., Mikoshiba, K., Gill, D. L., Han, S., and Wang, Y. (2020) Type 3 Inositol 1,4,5-Trisphosphate Receptor is a Crucial Regulator of Calcium Dynamics Mediated by Endoplasmic Reticulum in HEK Cells. Cells 9, 10.3390/cells9020275

7. Ando, H., Hirose, M., and Mikoshiba, K. (2018) Aberrant IP3 receptor activities revealed by comprehensive analysis of pathological mutations causing spinocerebellar ataxia 29. Proc Natl Acad Sci U S A 115, 12259–12264

8. Wang, Y. J., Huang, J., Liu, W., Kou, X., Tang, H., Wang, H., Yu, X., Gao, S., Ouyang, K., and Yang, H. T. (2017) IP3R-mediated Ca2+ signals govern hematopoietic and cardiac divergence of Flk1+ cells via the calcineurin-NFATc3-Etv2 pathway. Journal of molecular cell biology 9, 274–288

9. Ronkko, J., Rodriguez, Y., Rasila, T., Torregrosa-Munumer, R., Pennonen, J., Kvist, J., Kuuluvainen, E., Bosch, L. V. D., Hietakangas, V., Bultynck, G., Tyynismaa, H., and Ylikallio, E. (2023) Human IP(3) receptor triple knockout stem cells remain pluripotent despite altered mitochondrial metabolism. Cell Calcium 114, 102782

10. Wen, H., Xu, W. J., Jin, X., Oh, S., Phan, C. H., Song, J., Lee, S. K., and Park, S. (2015) The roles of IP3 receptor in energy metabolic pathways and reactive oxygen species homeostasis revealed by metabolomic and biochemical studies. Biochimica et biophysica acta 1853, 2937–2944

11. Lagos-Cabre, R., Ivanova, A., and Taylor, C. W. (2020) Ca(2+) Release by IP(3) Receptors Is Required to Orient the Mitotic Spindle. Cell reports 33, 108483

12. Young, M. P., Schug, Z. T., Booth, D. M., Yule, D. I., Mikoshiba, K., Hajnomicronczky, G., and Joseph, S. K. (2022) Metabolic adaptation to the chronic loss of Ca(2+) signaling induced by KO of IP(3) receptors or the mitochondrial Ca(2+) uniporter. J Biol Chem 298, 101436

13. Greenberg, M. E., Ziff, E. B., and Greene, L. A. (1986) Stimulation of neuronal acetylcholine receptors induces rapid gene transcription. Science 234, 80–83

14. Greenberg, M. E., and Ziff, E. B. (1984) Stimulation of 3T3 cells induces transcription of the c-fos proto-oncogene. Nature 311, 433–438

15. Thiel, G., Schmidt, T., and Rossler, O. G. (2021) Ca(2+) Microdomains, Calcineurin and the Regulation of Gene Transcription. Cells 10

16. West, A. E., Chen, W. G., Dalva, M. B., Dolmetsch, R. E., Kornhauser, J. M., Shaywitz, A. J., Takasu, M. A., Tao, X., and Greenberg, M. E. (2001) Calcium regulation of neuronal gene expression. Proc Natl Acad Sci U S A 98, 11024–11031

17. Masaki, T., and Shimada, M. (2022) Decoding the Phosphatase Code: Regulation of Cell Proliferation by Calcineurin. International journal of molecular sciences 23

18. Roche, E., and Prentki, M. (1994) Calcium regulation of immediate-early response genes. Cell Calcium 16, 331–338

19. Yeh, Y. C., and Parekh, A. B. (2018) CRAC Channels and Ca(2+)-Dependent Gene Expression. in Calcium Entry Channels in Non-Excitable Cells (Kozak, J. A., and Putney, J. W., Jr. eds.), Boca Raton (FL). pp 93–106

20. Luo, J., Busillo, J. M., and Benovic, J. L. (2008) M3 muscarinic acetylcholine receptor-mediated signaling is regulated by distinct mechanisms. Mol Pharmacol 74, 338–347

21. Arguin, G., Caron, A. Z., Elkoreh, G., Denault, J. B., and Guillemette, G. (2010) The transcription factors NFAT and CREB have different susceptibilities to the reduced Ca2+ responses caused by the knock down of inositol trisphosphate receptor in HEK 293A cells. Cellular physiology and biochemistry : international journal of experimental cellular physiology, biochemistry, and pharmacology 26, 629–640

22. Macian, F. (2005) NFAT proteins: key regulators of T-cell development and function. Nat Rev Immunol 5, 472–484

23. Bekker-Jensen, D. B., Kelstrup, C. D., Batth, T. S., Larsen, S. C., Haldrup, C., Bramsen, J. B., Sorensen, K. D., Hoyer, S., Orntoft, T. F., Andersen, C. L., Nielsen, M. L., and Olsen, J. V. (2017) An Optimized Shotgun Strategy for the Rapid Generation of Comprehensive Human Proteomes. Cell systems 4, 587–599 e584

24. Kar, P., and Parekh, A. B. (2015) Distinct spatial Ca2+ signatures selectively activate different NFAT transcription factor isoforms. Mol Cell 58, 232–243

25. Yoast, R. E., Emrich, S. M., Zhang, X., Xin, P., Johnson, M. T., Fike, A. J., Walter, V., Hempel, N., Yule, D. I., Sneyd, J., Gill, D. L., and Trebak, M. (2020) The native ORAI channel trio underlies the diversity of Ca(2+) signaling events. Nature communications 11, 2444

26. Hogan, P. G., Chen, L., Nardone, J., and Rao, A. (2003) Transcriptional regulation by calcium, calcineurin, and NFAT. Genes & development 17, 2205–2232

27. Macian, F., Lopez-Rodriguez, C., and Rao, A. (2001) Partners in transcription: NFAT and AP-1. Oncogene 20, 2476–2489

28. Steven, A., Friedrich, M., Jank, P., Heimer, N., Budczies, J., Denkert, C., and Seliger, B. (2020) What turns CREB on? And off? And why does it matter? Cellular and molecular life sciences : CMLS 77, 4049–4067

29. Dewenter, M., von der Lieth, A., Katus, H. A., and Backs, J. (2017) Calcium Signaling and Transcriptional Regulation in Cardiomyocytes. Circ Res 121, 1000–1020

30. Screaton, R. A., Conkright, M. D., Katoh, Y., Best, J. L., Canettieri, G., Jeffries, S., Guzman, E., Niessen, S., Yates, J. R., 3rd, Takemori, H., Okamoto, M., and Montminy, M. (2004) The CREB coactivator TORC2 functions as a calcium- and cAMP-sensitive coincidence detector. Cell 119, 61–74

31. Katoh, Y., Takemori, H., Lin, X. Z., Tamura, M., Muraoka, M., Satoh, T., Tsuchiya, Y., Min, L., Doi, J., Miyauchi, A., Witters, L. A., Nakamura, H., and Okamoto, M. (2006) Silencing the constitutive active transcription factor CREB by the LKB1-SIK signaling cascade. FEBS J 273, 2730–2748

32. Johannessen, M., Delghandi, M. P., and Moens, U. (2004) What turns CREB on? Cellular signalling 16, 1211–1227

33. Nixon, J. S., Bishop, J., Bradshaw, D., Davis, P. D., Hill, C. H., Elliott, L. H., Kumar, H., Lawton, G., Lewis, E. J., Mulqueen, M., and, et al. (1992) The design and biological properties of potent and selective inhibitors of protein kinase C. Biochem Soc Trans 20, 419–425

34. Rimessi, A., Rizzuto, R., and Pinton, P. (2007) Differential recruitment of PKC isoforms in HeLa cells during redox stress. Cell Stress Chaperones 12, 291–298

35. Kim, H., Na, Y. R., Kim, S. Y., and Yang, E. G. (2016) Protein Kinase C Isoforms Differentially Regulate Hypoxia-Inducible Factor-1alpha Accumulation in Cancer Cells. J Cell Biochem 117, 647–658

36. Karin, M., Liu, Z., and Zandi, E. (1997) AP-1 function and regulation. Current opinion in cell biology 9, 240–246

37. Nakabeppu, Y., and Nathans, D. (1991) A naturally occurring truncated form of FosB that inhibits Fos/Jun transcriptional activity. Cell 64, 751–759

38. Rosen, L. B., Ginty, D. D., Weber, M. J., and Greenberg, M. E. (1994) Membrane depolarization and calcium influx stimulate MEK and MAP kinase via activation of Ras. Neuron 12, 1207–1221

39. Agell, N., Bachs, O., Rocamora, N., and Villalonga, P. (2002) Modulation of the Ras/Raf/MEK/ERK pathway by Ca(2+), and calmodulin. Cellular signalling 14, 649–654

40. Chao, T. S., Foster, D. A., Rapp, U. R., and Rosner, M. R. (1994) Differential Raf requirement for activation of mitogen-activated protein kinase by growth factors, phorbol esters, and calcium. J Biol Chem 269, 7337–7341

41. Kupzig, S., Walker, S. A., and Cullen, P. J. (2005) The frequencies of calcium oscillations are optimized for efficient calcium-mediated activation of Ras and the ERK/MAPK cascade. Proc Natl Acad Sci U S A 102, 7577–7582

42. Enslen, H., Tokumitsu, H., Stork, P. J., Davis, R. J., and Soderling, T. R. (1996) Regulation of mitogen-activated protein kinases by a calcium/calmodulin-dependent protein kinase cascade. Proc Natl Acad Sci U S A 93, 10803–10808

43. Berry, C. T., May, M. J., and Freedman, B. D. (2018) STIM- and Orai-mediated calcium entry controls NF-kappaB activity and function in lymphocytes. Cell Calcium 74, 131–143

44. Peng, H. Y., Lucavs, J., Ballard, D., Das, J. K., Kumar, A., Wang, L., Ren, Y., Xiong, X., and Song, J. (2021) Metabolic Reprogramming and Reactive Oxygen Species in T Cell Immunity. Front Immunol 12, 652687

45. Vaeth, M., and Feske, S. (2018) NFAT control of immune function: New Frontiers for an Abiding Trooper. F1000Res 7, 260

46. Abate, C., Patel, L., Rauscher, F. J., 3rd, and Curran, T. (1990) Redox regulation of fos and jun DNA-binding activity in vitro. Science 249, 1157–1161

47. Morgan, M. J., and Liu, Z. G. (2011) Crosstalk of reactive oxygen species and NF-kappaB signaling. Cell research 21, 103–115

48. Ichiki, T., Tokunou, T., Fukuyama, K., Iino, N., Masuda, S., and Takeshita, A. (2003) Cyclic AMP response element-binding protein mediates reactive oxygen species-induced c-fos expression. Hypertension 42, 177–183

49. Aggarwal, V., Tuli, H. S., Varol, A., Thakral, F., Yerer, M. B., Sak, K., Varol, M., Jain, A., Khan, M. A., and Sethi, G. (2019) Role of Reactive Oxygen Species in Cancer Progression: Molecular Mechanisms and Recent Advancements. Biomolecules 9

50. Quinlan, C. L., Gerencser, A. A., Treberg, J. R., and Brand, M. D. (2011) The mechanism of superoxide production by the antimycin-inhibited mitochondrial Q-cycle. J Biol Chem 286, 31361–31372

51. Murphy, M. P., Bayir, H., Belousov, V., Chang, C. J., Davies, K. J. A., Davies, M. J., Dick, T. P., Finkel, T., Forman, H. J., Janssen-Heininger, Y., Gems, D., Kagan, V. E., Kalyanaraman, B., Larsson, N. G., Milne, G. L., Nystrom, T., Poulsen, H. E., Radi, R., Van Remmen, H., Schumacker, P. T., Thornalley, P. J., Toyokuni, S., Winterbourn, C. C., Yin, H., and Halliwell, B. (2022) Guidelines for measuring reactive oxygen species and oxidative damage in cells and in vivo. Nat Metab 4, 651–662

52. Gutscher, M., Pauleau, A. L., Marty, L., Brach, T., Wabnitz, G. H., Samstag, Y., Meyer, A. J., and Dick, T. P. (2008) Real-time imaging of the intracellular glutathione redox potential. Nature methods 5, 553–559

53. Pak, V. V., Ezerina, D., Lyublinskaya, O. G., Pedre, B., Tyurin-Kuzmin, P. A., Mishina, N. M., Thauvin, M., Young, D., Wahni, K., Martinez Gache, S. A., Demidovich, A. D., Ermakova, Y. G., Maslova, Y. D., Shokhina, A. G., Eroglu, E., Bilan, D. S., Bogeski, I., Michel, T., Vriz, S., Messens, J., and Belousov, V. V. (2020) Ultrasensitive Genetically Encoded Indicator for Hydrogen Peroxide Identifies Roles for the Oxidant in Cell Migration and Mitochondrial Function. Cell Metab 31, 642–653 e646

54. Ma, Q. (2013) Role of nrf2 in oxidative stress and toxicity. Annu Rev Pharmacol Toxicol 53, 401–426

55. Kuwahara, K., Wang, Y., McAnally, J., Richardson, J. A., Bassel-Duby, R., Hill, J. A., and Olson, E. N. (2006) TRPC6 fulfills a calcineurin signaling circuit during pathologic cardiac remodeling. J Clin Invest 116, 3114–3126

56. Katona, M., Bartok, A., Nichtova, Z., Csordas, G., Berezhnaya, E., Weaver, D., Ghosh, A., Varnai, P., Yule, D. I., and Hajnoczky, G. (2022) Capture at the ER-mitochondrial contacts licenses IP(3) receptors to stimulate local Ca(2+) transfer and oxidative metabolism. Nature communications 13, 6779

57. Rouillard, A. D., Gundersen, G. W., Fernandez, N. F., Wang, Z., Monteiro, C. D., McDermott, M. G., and Ma’ayan, A. (2016) The harmonizome: a collection of processed datasets gathered to serve and mine knowledge about genes and proteins. Database (Oxford) 2016

58. Pont, J. N., McArdle, C. A., and Lopez Bernal, A. (2012) Oxytocin-stimulated NFAT transcriptional activation in human myometrial cells. Mol Endocrinol 26, 1743–1756

59. Demozay, D., Tsunekawa, S., Briaud, I., Shah, R., and Rhodes, C. J. (2011) Specific glucose-induced control of insulin receptor substrate-2 expression is mediated via Ca2+-dependent calcineurin/NFAT signaling in primary pancreatic islet beta-cells. Diabetes 60, 2892–2902

60. Naito, S., Shimizu, S., Maeda, S., Wang, J., Paul, R., and Fagin, J. A. (1998) Ets-1 is an early response gene activated by ET-1 and PDGF-BB in vascular smooth muscle cells. Am J Physiol 274, C472–480

61. Tsao, H. W., Tai, T. S., Tseng, W., Chang, H. H., Grenningloh, R., Miaw, S. C., and Ho, I. C. (2013) Ets-1 facilitates nuclear entry of NFAT proteins and their recruitment to the IL-2 promoter. Proc Natl Acad Sci U S A 110, 15776–15781

62. Zagranichnaya, T. K., Wu, X., Danos, A. M., and Villereal, M. L. (2005) Gene expression profiles in HEK-293 cells with low or high store-operated calcium entry: can regulatory as well as regulated genes be identified? Physiol Genomics 21, 14–33

63. Dolmetsch, R. E., Xu, K., and Lewis, R. S. (1998) Calcium oscillations increase the efficiency and specificity of gene expression. Nature 392, 933–936

64. Li, W., Llopis, J., Whitney, M., Zlokarnik, G., and Tsien, R. Y. (1998) Cell-permeant caged InsP3 ester shows that Ca2+ spike frequency can optimize gene expression. Nature 392, 936–941

65. Lenz, J. C., Reusch, H. P., Albrecht, N., Schultz, G., and Schaefer, M. (2002) Ca2+-controlled competitive diacylglycerol binding of protein kinase C isoenzymes in living cells. J.Cell Biol. 159, 291–302

66. Kotla, S., Singh, N. K., Kirchhofer, D., and Rao, G. N. (2017) Heterodimers of the transcriptional factors NFATc3 and FosB mediate tissue factor expression for 15(S)-hydroxyeicosatetraenoic acid-induced monocyte trafficking. J Biol Chem 292, 14885–14901

67. Hasan, G. (2013) Intracellular signaling in neurons: unraveling specificity, compensatory mechanisms and essential gene function. Curr Opin Neurobiol 23, 62–67

68. Robinson, M. D., McCarthy, D. J., and Smyth, G. K. (2010) edgeR: a Bioconductor package for differential expression analysis of digital gene expression data. Bioinformatics 26, 139–140

69. Hortenhuber, M., Toledo, E. M., Smedler, E., Arenas, E., Malmersjo, S., Louhivuori, L., and Uhlen, P. (2017) Mapping genes for calcium signaling and their associated human genetic disorders. Bioinformatics 33, 2547–2554

70. Lambert, S. A., Jolma, A., Campitelli, L. F., Das, P. K., Yin, Y., Albu, M., Chen, X., Taipale, J., Hughes, T. R., and Weirauch, M. T. (2018) The Human Transcription Factors. Cell 172, 650–665

71. Rath, S., Sharma, R., Gupta, R., Ast, T., Chan, C., Durham, T. J., Goodman, R. P., Grabarek, Z., Haas, M. E., Hung, W. H. W., Joshi, P. R., Jourdain, A. A., Kim, S. H., Kotrys, A. V., Lam, S. S., McCoy, J. G., Meisel, J. D., Miranda, M., Panda, A., Patgiri, A., Rogers, R., Sadre, S., Shah, H., Skinner, O. S., To, T. L., Walker, M. A., Wang, H., Ward, P. S., Wengrod, J., Yuan, C. C., Calvo, S. E., and Mootha, V. K. (2021) MitoCarta3.0: an updated mitochondrial proteome now with sub-organelle localization and pathway annotations. Nucleic acids research 49, D1541–D1547

72. Subramanian, A., Tamayo, P., Mootha, V. K., Mukherjee, S., Ebert, B. L., Gillette, M. A., Paulovich, A., Pomeroy, S. L., Golub, T. R., Lander, E. S., and Mesirov, J. P. (2005) Gene set enrichment analysis: a knowledge-based approach for interpreting genome-wide expression profiles. Proc Natl Acad Sci U S A 102, 15545–15550

